# The trembling hand unraveled: motor and valuation elements in the neural sources of choice inconsistency

**DOI:** 10.1101/2022.12.20.521216

**Authors:** Vered Kurtz-David, Asaf Madar, Adam Hakim, Noa Palmon, Dino J Levy

## Abstract

Extensive evidence shows that humans are inconsistent with their choices. Yet, the neural mechanism underlying this type of choices remains unknown. Here, we aim to show that inconsistent choice is tied to the valuation process, but can also arise from motor errors during task execution. We report the results from three behavioral and neuroimaging studies. Subjects completed a risky-choice task to test their inconsistency levels, followed by two novel tasks, explicitly designed to examine motor output. We recorded mouse trajectories during task execution and designed 34 features to analyze motor dynamics in an exploratory manner. We show that motor dynamics predict inconsistency levels, even when motor output was absent any valuation elements. In the neuroimaging study, we show that inconsistency is associated with value brain circuits, but at the same time, is also related to activity in motor circuits. These findings suggest that (at least) two neural sources of noise contribute to inconsistent choice behavior.

## Introduction

Imagine you are in the arcade, trying to pull your favorite blue teddy bear out of the claw machine. However, for some reason, you end up grabbing a less preferred red teddy bear instead of the blue one. This inconsistency between your “true” preference and your actual choice can be the result of (at least) two different kinds of errors: it might be that you miscalculated the values of each teddy bear, leading to suboptimal inference^1,2^. It might also be that you accurately calculated the “correct” values of the options, but had a moment of motor "clumsiness", when you reached out to grab the stuffed animal.

In the 1970s, Nobel-prize laureate Reinhard Selten has coined the term “*trembling hand error*”, referring to the possibility that decision makers choose suboptimal options with small probabilities^3^. This notion has become a synonym for accidental outcomes in decision-making^4–7^. Nonetheless, as the term implies, and demonstrated in the arcade example, such mistakes could be the result of altered value representations, and/or due to motor tremor that accompanied choice execution.

Value-based choice is guided by common currency valuations of options in value-related brain regions^8,9^. Previous studies demonstrated how explore-exploit strategies^10,11^ or recent reward history^12^ can lead to stochastic decision-making and suboptimal value-based choices.

Others argued for an additional source for these behaviors – random neural activity or noise^1,2,11,13–15^, which can either arise in sensory regions during the perception of choice options^16^, or in decision circuits of the value network^17^, sometimes referred to as inference noise^1^.

Importantly, the valuation of choice options is also reflected in the motor system through the selection of strategies in task execution like the latency of movement onset or its speed^18,19^. As such, it is prone to imprecisions originating from noise stemming from motor neural circuits^20,21^. While stochastic motor computations have been shown to affect performance in classic goal-directed motor tasks such as moving or grabbing of objects^20,22,23^, a lot less is known on how motor noise – which is independent of inference or sensory noise – contributes to value-based decision-making in general, and to inconsistency in choices in particular.

Choice inconsistency violates fundamental axioms in economic theory^24–26^, as it disobeys the law of transitivity and creates choice cycles. Inconsistency is considered as an irrational behavior, as it nullifies the feasibility of a preference-ordering utility construct^24^. Importantly, choice inconsistency occurs even when the choice environment remains unchanged, with no context manipulations induced by the experimenter^27–29^. As such, it resembles well-documented findings in psychophysics that noisy random mechanisms vary perception and motor output, even when the stimuli, environment, or task remain constant^16^. This suggests that choice inconsistency may stem from noisy neural processes in various brain systems.

In the current work, we aim to differentiate between two sources of neural stochasticity and argue that both sources contribute to choice inconsistency. The first source is related to neural activity in value-related brain areas (inference noise or value miscalculations), corresponding to a previous work from our lab^27^. The second source is related to noise during task execution and originates from motor-related brain regions. Demonstrating the contribution of motor-related noise to violations of economic theory will provide scholars with a more mechanistic and neural-based approach to choice inconsistency than the general cognitive theories that dominate the literature^30^.

To this aim, we use a well-studied task with a simple graphical interface^31^ for eliciting subjects’ inconsistency levels based on choices from linear budget sets^27–29,31,32^. We leverage the vast dynamic information concealed within the process of executing each choice in our task. Therefore, in addition to the analysis of choice behavior, we also study the dynamics of the motor movements leading up to choices via the analysis of mouse trajectories. Crucially though, the greatest challenge when attempting to tease apart the two sources for suboptimality in choice, is to disentangle motor errors in task execution from value miscalculations within the decision process per se. To overcome this, we present a novel task design, which solely captures motor output free of any value computations. We then use the rich dynamic information of the mouse tracking data, to quantify the contribution of each of the two sources to the intensity of inconsistency levels.

Previous works that examined the role of motor computations in value-based decision-making have showed that motor output can alter preferenece-ordering^33,34^, or pointed at mutual learning mechanisms across the two domains^35^. Interestingly, recent studies that have aimed to quantify the role of motor computations on stochasticity in value-based choice, only found a negligible effect^1,11^. Nevertheless, these works have employed motor paradigms which were too simplistic to trace such an effect. That is to say, we pose that motor stochasticity plays a substantial role in any value-based decision process, but to fully test it, one would require a sufficiently-complex motor execution embedded into the task design. Here, we aim to quantify, on a trial-by-trial basis, the magnitude of motor-related computations in inconsistent choices, when a substantial motor element is required for task execution. We then directly link it to its corresponding neural footprints in motor brain regions.

Our work leverages the vast literature in recent years on mouse tracking tools, usually used in binary choice tasks, to investigate cognitive processes underlying subjects’ behavior in general^36,37^, and value-based decision-making mechanisms in particular^38–40^. Importantly, not only does the mouse tracking data allow us to differentiate the sources for suboptimal choice, but it also enables us to identify the particular elements within the motor movement that contribute to inconsistent choice behavior. Our mouse tracking analysis strategy thus takes an exploratory model-free approach, aiming to identify multiple aspects of movement that could be tied to inconsistency. Finally, we employ neuroimaging to examine the neural footprints of motor dynamics, and to pinpoint the different neural circuits related to the two sources that contribute to inconsistent choices.

We report the results from eight tasks in three separate studies: a behavioral study (n=89), a neuroimaging study (n=42), and a replication of the behavioral study (n=69). The replication study aimed to validate our findings due to the exploratory nature of our analysis approach. Our main behavioral findings show that motor output, even when absent value modulation, was able to predict inconsistency levels, and thus, can be regarded as a source for suboptimality in choice. In the neuroimaging study, we replicate our previous findings, and show that inconsistent choice is related to noisy neural networks during the computation of value^27^, but at the same time, we also demonstrate that inconsistency is strongly associated with activations in motor brain regions. Taken together, these findings suggest a substantial role of motor circuits in inconsistent choice.

## Results

### Inconsistency and choice behavior

Subjects completed a well-validated risky choice task to test their inconsistency levels^27,28^ (Fig. 1a left panel and Methods). On each trial, subjects were facing 50-50% lotteries between the X and Y amounts, and had to choose their most desired lottery. The set of all available lotteries was represented along a *budget line*. For example, the (53,18) coordinate (red dot) on the line presented in Fig 1a is a lottery with 50% chance to win 53 tokens, and 50% chance to win 18 tokens. The slope of the line indicated the substitution (price ratio) between the X and Y amounts, whereas the distance of the line from the axes-origin indicated the total token-amount to be won (*endowment*). Slopes and endowments were varied across trials. Subjects used a mouse (behavioral and replication studies) or a trackball (neuroimaging study) to choose their most desired lottery, while their mouse trajectories were recorded.

**Fig 1.**
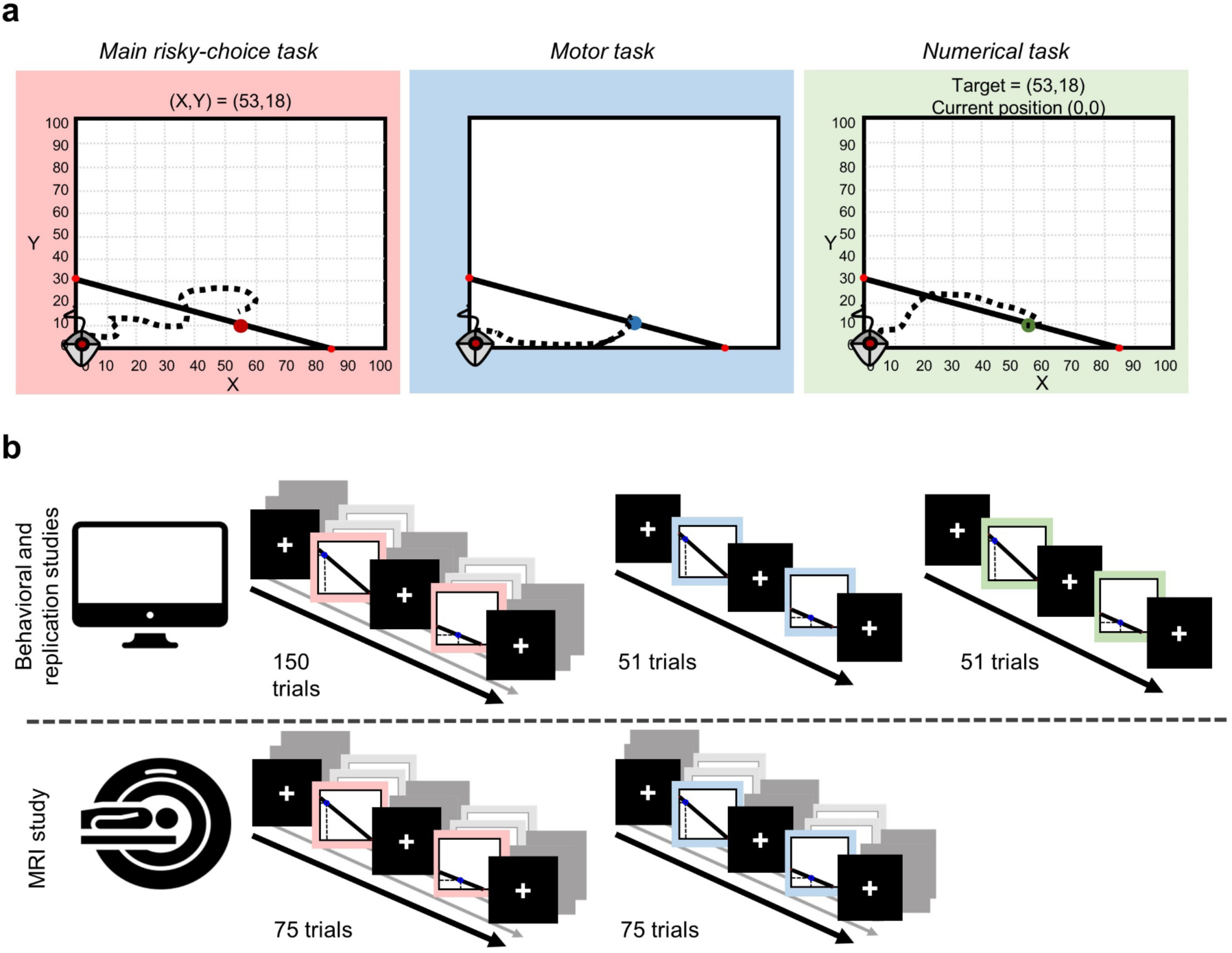
Experimental design. (a) In all tasks, at the beginning of each trial we set the mouse cursor at the axes-origin, and recorded mouse trajectories during the trial. In the behavioral and replication studies subjects used a mouse while in the fMRI study they used a trackball to submit their choices. *Main risky-choice task (left).* Subjects were presented with a budget line with 50/50 lotteries between two accounts, labeled *X* and *Y*. Each point on the budget line represented a different lottery between the *X* and *Y* accounts. Subjects had to choose their preferred lottery (a bundle of the *X* and *Y* accounts) out of all possible lotteries along the line. For example, the bundle (53,18, red dot) corresponds to a lottery with a 50% chance to win 53 tokens (account *X*), and a 50% chance to win 18 tokens (account *Y*). Each trial presented a different budget line with a different slope (the substitution, or price ratio, between *X* and *Y*) and endowment (the distance from the axes-origin), which were randomized across trials. *Middle and right panels.* Non-value tasks, aimed to study motor output, absent value-based decision-making. The lines and predefined targets were the budget lines and actual choices from the main task, respectively. *Motor task* (middle, used in both the behavioral and neuroimaging studies). Subjects had to move the cursor to the predefined blue target. Note, that no numerical information was presented, therefore not available for subjects to guide their choices. *Numerical task (right,* used solely in the behavioral study). Subjects had to reach the predefined {*X*, *Y*} target coordinates. The title displayed subjects’ current position (also see Methods). (b) Experimental procedures. *Upper panel* - in the behavioral and replication studies (n=89 and n=69, respectively), subjects completed 150 trials of the main task divided to 3 blocks of 50 trials, followed by one block of 51 trials in each of the non-value tasks (counter-balanced across subjects). *Bottom panel –* in the neuroimaging study (n=42), subjects completed 75 trials in the main and in the motor tasks, each divided to 3 blocks of 25 trials. We thus report results from all eight tasks across the three studies.

Subjects had no difficulty understanding the task (Supplementary Fig. 1) and responded to changes in prices (slopes), following the Law of Demand (Fig. 2b). Based on subjects’ choices in the risky choice task, we computed their subject-by-subject number of violations of the General Axiom of Revealed Preference (GARP)^24^. We report two different subject-by-subject inconsistency measures for the intensity of violations: the non-parametric Afriat Index (AI)^41^ and the parametric Money Metric Index (MMI)^32^. See Methods for a brief intuition on how to interpret the indices, Supplementary Note 3 for a more detailed account, and Supplementary Fig. 7 for a graphical illustration of the indices. Overall, subjects exhibited a large heterogeneity in their inconsistency levels (see Fig. 2a and Supplementary Fig. 8 for inter-indices correlations).

**Fig 2.**
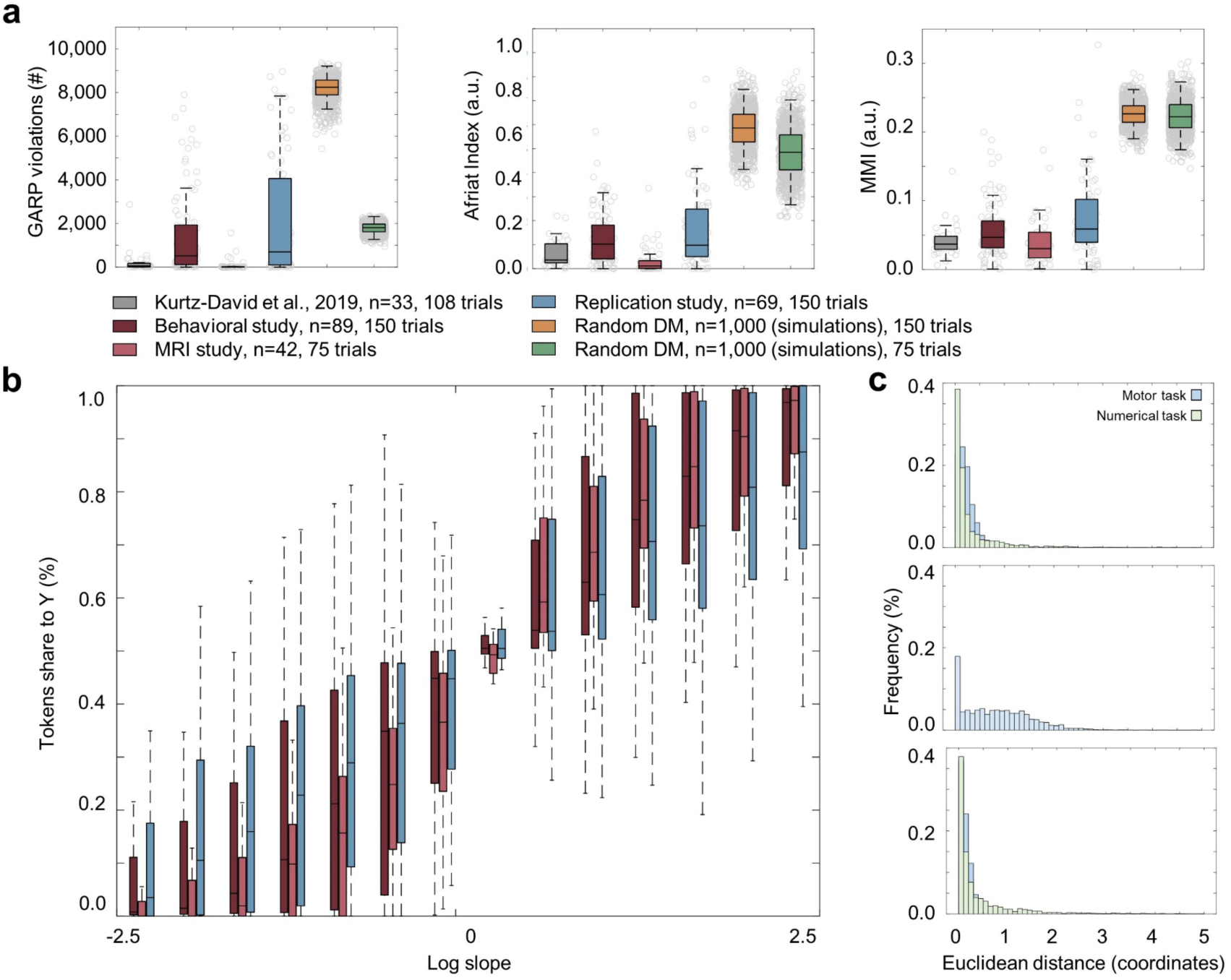
Inconsistency levels and imprecision in the non-value motor tasks. (a) Distributions of inconsistency indices among subjects in the three studies, compared to subjects in our previous study^27^ and to 1,000 simulated random decision-makers. In the behavioral study (red bars), only 3 subjects (out of 89, 3.4%) were fully consistent with GARP. Subjects made on average 1,509.1 GARP violations (min=0, max=8,258, std = 2,036.0). Subjects had an average AI score of 0.128 (min=0, max=0.572, std=0.113) and an MMI score of 0.058 (min=0.001, max=0.200, std=0.042). In the neuroimaging study (pink bars), 8 subjects (out of 42, 19.0%) were fully consistent with GARP. Subjects made on average 91.0 GARP violations (min=0, max=1,568, std = 289.5) with an average AI score of 0.037 (min=0, max=0.336, std=0.006) and an MMI score of 0.040 (min=0.001, max=0.174, std=0.036). In the replication study, 3 subjects (out of 69, 4.3%) were fully consistent with GARP. Subjects made on average 2,101.6 GARP violations (std=2,742.3, min=0, max=8,962) with a mean AI=0.1702 (std=0.1774, min=0, max=0.7895) and MMI=0.0756 (std=0.069, min=0.0001, max=0.3267). Whiskers indicate one std. (b) The share of total endowment (expenditure) subjects allocated to the Y account as a function of the price ratio 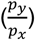, the slope of the budget set, increased as the Y account became cheaper. Price ratios are grouped into 15 bins. In each boxplot center line indicates the median share of tokens; box limits indicate upper and lower quartiles; and whiskers indicate one std. (c) Distributions of Euclidean dist. (in coordinate units) between target coordinates and the coordinates of mouse-button presses in the non-value tasks. Euclidean dist. could range between 0 and 141.42 (representing the longest diagonal within the axes’ grid). *Top -* Behavioral study. Motor task – mean: 0.597, std: 3.403, min: 0, max: 75.479; Numerical task – mean: 0.784, std: 4.670, min: 0, max: 116.885; *middle -* Neuroimaging study: motor task *-* mean: 0.849, std: 0.795, min: 0, max: 14.677. *Bottom –* replication study. Motor task – mean: 1.258, std=7.07, min=0, max=95.65; Numerical task – mean=0.886, std=5.750, min=0, max=128.7. Distributions of Euclidean distances are highly asymmetrical and skewed at 0, suggesting very high accuracy rates and similar button-press coordinates across tasks. Note that the Euclidean distances in the neuroimaging study (middle panel) are higher than in the behavioral study (top panel, p<0.0001, two-sample, one-sided Wilcoxon rank-sum test), probably due to some difficulty to maneuver the trackball inside the MRI scanner.

The number of violations is heavily dependent on the number of trials that subjects encountered^42^. Hence, because subjects in the three samples in our study completed a different number of trials (behavioral and replication studies: 150 trials, neuroimaging study: 75 trials), we do not compare subjects’ inconsistency levels across the different samples. Instead, we compare their choices with 1,000 simulated random decision-makers^42^, to show that even though subjects were inconsistent in their choices, they did not choose at random. Indeed, across all three inconsistency indices under examination, inconsistency levels among participants in our study were substantially lower than those of the simulated choosers (Behavioral study: GARP violations: z=15.5244, q(FDR)<0.0001; AI: z=15.4744, q(FDR)<0.0001; MMI: z=15.6188, q(FDR)<0.0001. n_1_=89, n_2_=1,000. Neuroimaging study: GARP violations: z=10.8915, q(FDR)<0.0001; AI: z=10.9532; q(FDR)<0.0001; MMI: z=10.9788, q(FDR)<0.0001. n_1_=42, n_2_=1,000. Replication study: GARP violations: z=12.7769, q(FDR)<0.0001; AI: z=12.4674, q(FDR)<0.0001; MMI: z=13.000, q(FDR)<0.0001. n_1_=69, n_2_=1,000. One-sided two-sample Wilcoxon rank sum test, Fig. 2a, yellow and green boxplots). Importantly, in the absence of an appropriate statistical measure, Bronars^42^ suggested to use the frequency of simulated choosers who violated GARP as a measure for the power of the inconsistency test. As none of the simulated choosers complied with GARP, we conclude that in all three studies, our task had a power of (at least) 99.999% for detecting (in)consistencies in subjects’ choices.

The rest of the paper will focus on trial-by-trial analyses. Similarly to our previous study^27^, for the trial-level analysis we employ a Leave-one-out procedure on MMI to yield the trial-level-MMI index (henceforth: MMI, Methods and Supplementary Note 3).

### Extracting motor related features

As a first step towards understanding motor output in our task, after analyzing subjects’ submitted choices and measuring their level of choice inconsistency in each trial, we systematically analyzed the dynamics leading up to a choice (henceforth, *“choice dynamics”*). Therefore, in all three studies we recorded the mouse trajectories of motion pathways until the final choice was made (see Supplementary Note 4 and Supplementary Fig. 2 for descriptive results from the mouse tracking analysis). To take full advantage of the information concealed in the rich datasets provided by the mouse trajectories, we extracted various features from the mouse trajectories.

Our aim was to characterize as many aspects of the choice dynamics as possible and to relate them to choice inconsistencies. We thus extracted a total of 34 different mouse features, such as the mean velocity of movement (meanVel), layover time at the axes-origin (Layover), the aggregated curvature of the trajectory (Curveness), time spent outside the axes-grid (XTimeOutofBounds and YTimeOutofBounds) or the entropy of the trajectory pathway (XSampEn4 and YSampEn4). Most of the extracted features (25 features out of 34) were based on standard features from the mouse tracking literature^36,43–46^, while nine features were based on our own design (see Methods and Supplementary Note 2 for details). Our features design approach thus used well-validated features, but at the same time, also explored novel task-related features. Our goal was to use the high dimensional space created by the features to improve prediction results. The large number of features was therefore the main reason for running the replication study, which was aimed to strengthen our conclusions and to reduce any confounds that may had been created simply due to our choice of specific mouse features.

Fig. 3a-d visualize four representative features, Supplementary Table 3 provides a detailed list of all the features, and Supplementary Figs. 3-4 present the distributions of features in the behavioral and neuroimaging studies. Fig. 3e illustrates the distribution across subjects of mouse features’ values across trials (same features as in Fig. 3a-d) and suggests that there was high variability in response strategies within and across subjects.

**Fig 3.**
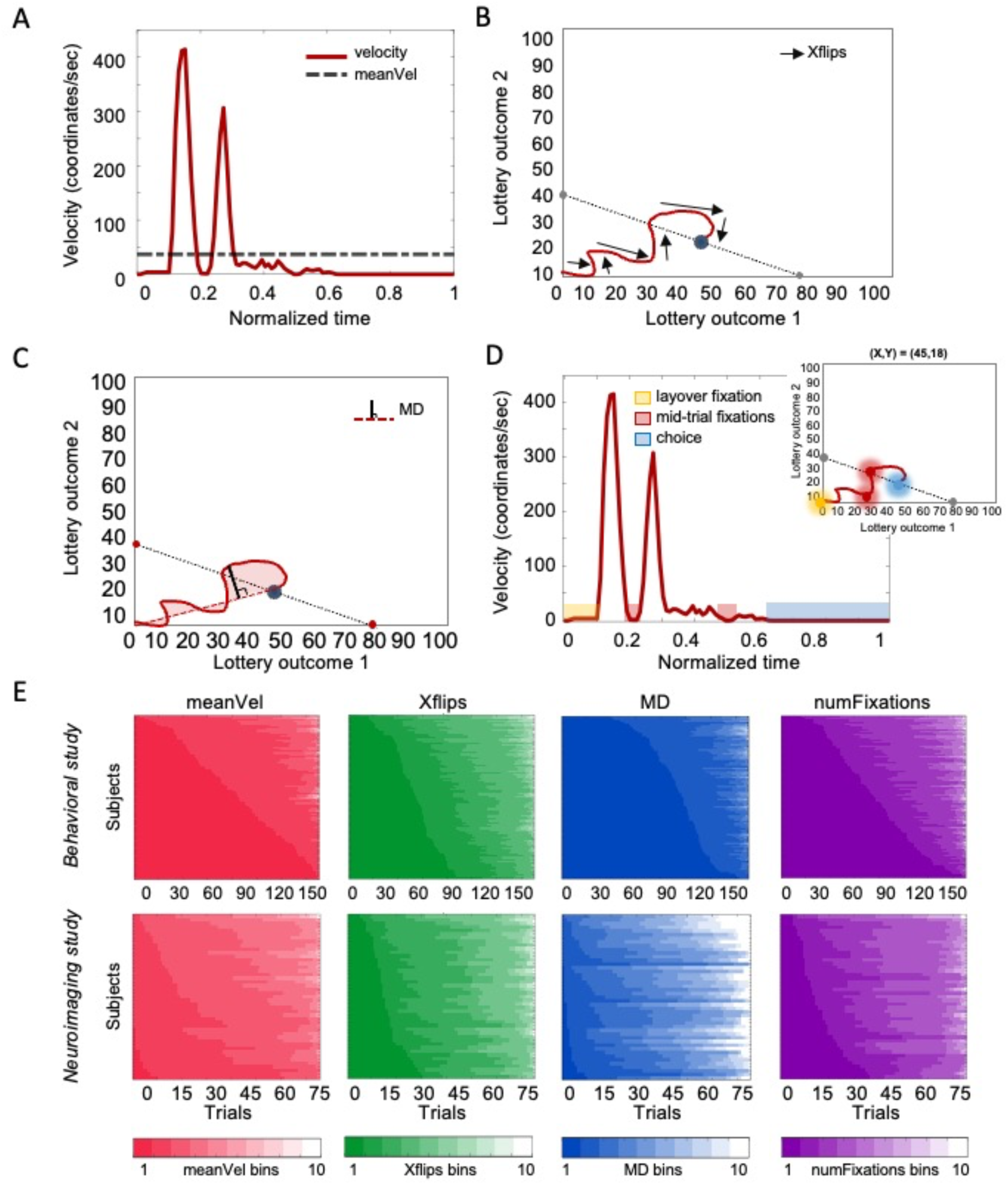
Mouse features. (a-d) Representative features (See Methods, Supplementary Table 3 and Supplementary Note 2). (a) *meanVel* – for each trial we measured the average velocity (dashed line) in coordinates/second (same trial as in Supplementary Fig. 2a, left panel). (b) *Xflips* – in each trial, we measured the number of changes in movement direction (flips) along the x-axis. (c) *MD* – in each trial we calculated the maximal distance between the actual mouse trajectory and the “choice line” – the shortest line connecting the axes origin and choice location. (d) *numFixations*– in each trial, we divided the graph to a 20x20 grid, such that each bin had a width of 5 tokens. Any bin in which the subject spent more than 0.2 seconds was considered a “mouse fixation”, meaning segments in which the subject ceased motion, excluding the actual choice. In the numFixations feature, we counted the number of such segments. Figure inset – spatial illustration of mouse fixations. (e) Distributions of trials based on features’ values (same features as in a-d). We grouped features into ten equally-spaced bins. Subjects are sorted by ascending order according to their share of trials classified to bin 1 (lowest values). Note that within subject (each row), trials are sorted by bins in an ascending order of feature-value, and not by chronological trial-order. (*Up* – features from the behavioral study, *Bottom* – features from the neuroimaging study).

### Mouse features from the risky-choice task predict inconsistency levels

Our main goal was to demonstrate that, in addition to value miscalculations (inference noise), choice inconsistency was influenced by the dynamics of motor characteristics originating in noisy neural computations in motor networks unrelated to value-based computations. As a first step towards this goal, we aimed to demonstrate that specific elements in motor output, captured by the extracted mouse features, could be related to subjects’ inconsistent choice behavior.

Since the mouse features were highly correlated (Fig. 4a) and to avoid overfitting, we induced a dimensionality reduction procedure on the mouse features. We conducted a linear regression with elastic net regularization (see Methods) that resulted in a subset of mouse features out of the total 34 features. The advantage of using this approach is that it maintains the original mouse features, which allows interpretability of the results. Specifically, 18 (behavioral study),11 (neuroimaging study) and 21 (replication study) features were selected by the elastic net regularization and significantly correlated with inconsistency levels (p<0.05, multiple linear model with subjects fixed effects, Fig. 4c, Supplementary Table 4, and Methods). This suggests that specific elements in task execution accounted for inconsistency levels.

**Fig 4.**
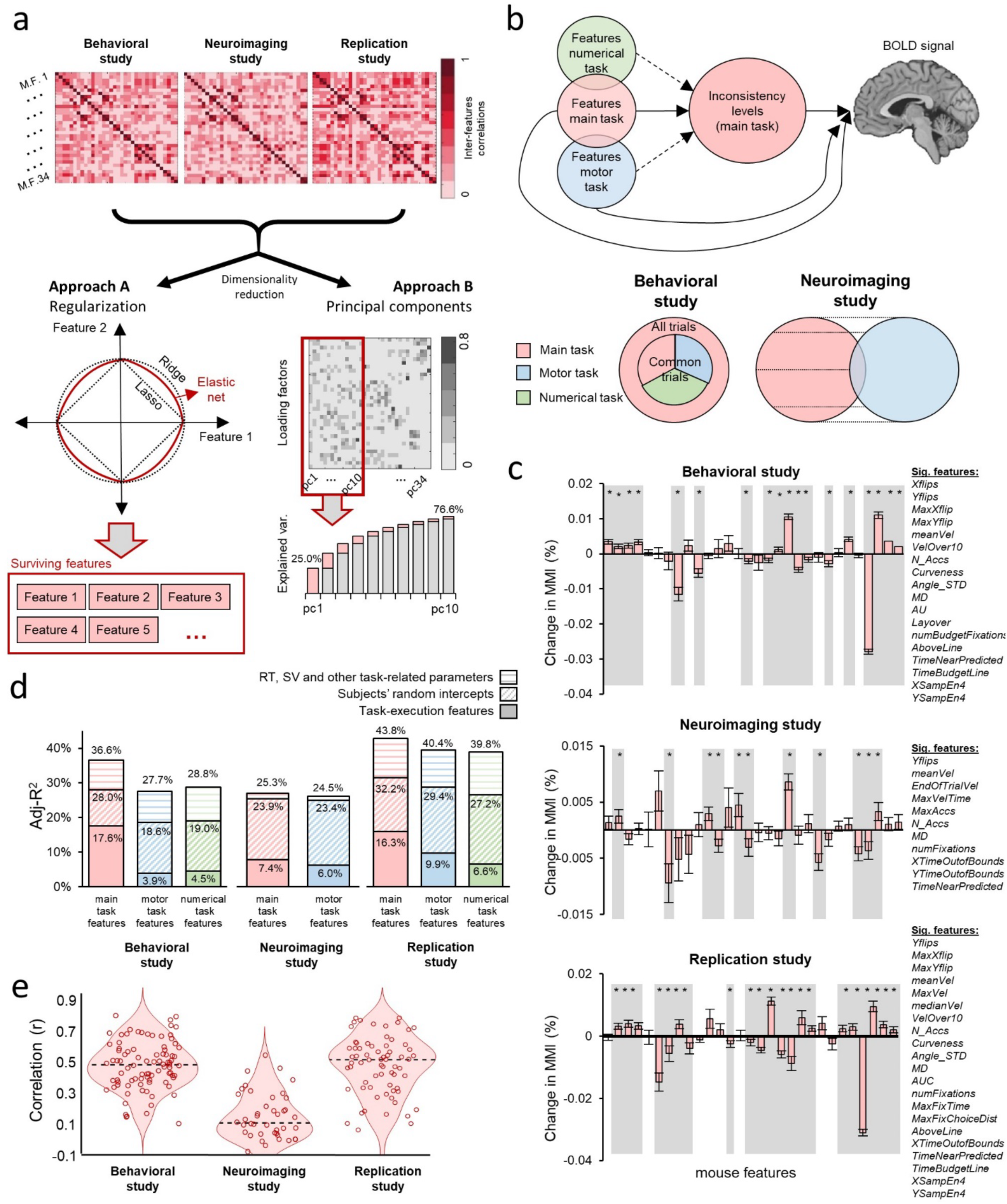
Mouse features and inconsistency levels. (a-b) Analysis strategy. (a) We took two dimensionality reduction approaches, in each of the studies, due to high dimensionality relative to the number of trials per subject, as well as due to inter-features correlations (upper row). In 84 (behavioral study), 56 (neuroimaging study) and 114 (replication study) pairs out of the 561 off-diagonal correlations, the mean correlation coefficient was higher or equal to 0.4 in absolute value. Approach A (*left)*: elastic net, which combines two methods for regularized regression (LASSO and Ridge) and maintains interpretability of the features. See Supplementary Table 4 for the list of features, which survived the elastic net regularization. Approach B (*right*): PCA. This approach preserves all features’ data on the expense of interpretability. In all further analyses, we used the first 10 components, which accounted for 76.6% (behavioral study), 77.3% (neuroimaging study) and 78.2% (replication study) of the variance in the data. (b) In the main risky-choice task, we used the selected features from Approach A and the PCs from Approach B to account for MMI scores. We then repeated the analysis in Approaches A and B with mouse features from the non-value motor tasks as out-of-sample predictors for MMI scores estimated in the value-based task. In the out-of-sample analysis, we solely used the common trials across tasks (51 trials in the behavioral and replication studies, and all 75 trials in the neuroimaging study). For the fMRI analysis, we used the selected features and looked for brain regions that correlated with both mouse features and MMI scores. (c) Mouse features in the main risky-choice task correlated with inconsistency levels in both studies (Approach A). Shaded areas indicate a significant predictor (p<0.05) in a subjects fixed-effects regression (see Methods). *Upper*: behavioral study. *Middle:* neuroimaging study. Bottom: replication study. (d) Adj-R^2^ from the analyses described in (b) (Approach A, selected features). Solid fill indicates regressions, which only included task-execution mouse features. Diagonal striped fill indicates subjects’ random intercept. Horizontal striped fill indicates additional explained variance from RT, SV, and other task-related parameters (see Methods). (e) Leave-one-subject-out prediction. Distribution of correlation coefficients between predicted inconsistency levels based on the selected mouse features from the main task and actual MMI scores. *Left -* Behavioral study: median: 0.491, min=0.117, max=0.805, std=0.153. *Middle:* neuroimaging study: median: 0.1177, min=-0.1789 max=0.5830, std=0.1652. *Right*: replication study: median: 0.514, min=-0.114, max=0.784, std=0.196. Dashed black line indicates median. Scatter shows individual correlation coefficients (behavioral study: N=89, neuroimaging study: N=42, replication study: N=69.). (c-e) See Supplementary Fig. 5 for the results for *Approach B: PCA*.

Goodness-of-fit measures (adj-R^2^) show that task-execution mouse features accounted for 17.6% (behavioral study), 7.4% (neuroimaging study) and 16.2% (replication study) of the variance of inconsistency scores (28.0%, 23.9% and 32.2%, once including random intercepts by subjects into the model, respectively. See Fig. 4d. See model equation in Methods). Note that this effect is achieved even when controlling for other constructs that can also be tied to inconsistency levels, such as reaction time (RT), the subjective value of the chosen lottery (SV), choice difficulty, and budget set parameters (horizontal striped bars in Fig. 4d, see Methods).

To further strengthen this finding, we conducted a leave-one-subject-out prediction, in which dimensionality reduction and model coefficients were trained on all trials from n-1 subjects and tested on the trials of the n^th^ subject (see Methods). We found a positive correlation between predicted and actual inconsistency levels for all subjects in the behavioral study, for 37 of 42 subjects in the neuroimaging study, and 68 of 69 subjects in the replication study (p<0.0001, Behavioral study: t(88)= 23.2446, median correlation r=0.4910. Neuroimaging study t(41)= 6.210, median correlation r=0.1177. Replication study: t(68)=19.896, median correlation r=0.5142. One-sided one-sample t-test, Fig. 4e). These results show that, indeed, choice dynamics in the value-based risk task were related to inconsistency levels, both in- and out-of-sample.

One may argue that the regularization provided by the elastic net approach is not sufficient to fully remove the collinearity between features. Thus, to check the robustness of our results, we also employed an unsupervised algorithm that reduced the dimensionality of the data, while simultaneously removing collinearities between the features. To this end, we conducted a principal component analysis (PCA; see Supplementary Note 5) on the mouse features. Importantly, this method preserved all the mouse features (as opposed to the elastic net), but at the expense of the interpretability of the components. We then used the results from the PCA and examined whether the principal components correlated with inconsistency levels in a subjects fixed-effects regression (Fig. 4b and Supplementary Fig. 5a). The PCA strategy yielded very similar results to the elastic-net approach (Supplementary Note 5 and Supplementary Fig. 5b-c).

To conclude, two analysis approaches, across three independent samples, indicated that motor components during task execution were related to subjects’ inconsistent choice behavior on a trial-by-trial level. However, mouse features from the risky-choice task did not solely capture “pure” motor computations, since subjects might have been actively engaging in value computations during the motor trajectory towards the final choice^47^. Accordingly, one may argue that the results presented in this section captured a combination of the valuation component of inconsistency, and motor components, which originated in motor circuits, hence only partially demonstrating our main goal. The next set of analyses addresses exactly this issue.

### Basic motor traits, unrelated to value modulation, also predict choice inconsistency

Since we aimed to demonstrate that inconsistency was influenced by the dynamics of motor characteristics originating in noisy computations in neural motor networks unrelated to value-based computations, we wanted to investigate subjects’ motor output outside the value-based choice framework. Therefore, following the main risky-choice task, subjects also completed novel non-value tasks (the *Motor task* and the *Numerical task*, Fig. 1a-b, Methods and Supplementary Notes 1). We designed the non-value tasks in such a manner that precluded any sort of a valuation process, yet would yield similar trajectories to the ones in the main risky-choice task. This design allowed us to relate task execution in the non-value tasks to inconsistency levels in the main risky-choice tasks, which could directly indicate on additional sources for choice inconsistency other than miscalculations of value. Here, too, we recorded mouse trajectories during task execution and extracted the same 34 mouse features that we extracted in the main risky-choice task (Fig. 3, Methods and Supplementary Note 2). To differentiate mouse features extracted from the risky-choice task (*choice dynamics)* from mouse features extracted from the non-value tasks, we refer the latter as “*motor dynamics”*.

In the *Motor task,* subjects had to reach a predefined circular target marked on a linear graph, similar to the linear budget set, but the graph had no numbers on the axes-grid, echoing well-known target-reaching tasks^48^. In the *Numerical task* subjects had to reach a predefined numerical target coordinate that lied on a linear graph. Here, subjects had to constantly determine whether their current location matched the abstract target they were given, thus resembling spatial navigation tasks. Importantly, the linear graphs in both non-value tasks were the actual budget lines from the main value task and the location of the predefined targets were identical to the location of the choices each subject made in the main choice task (Fig. 1a middle and right panels).

Mouse trajectories from the non-value motor tasks were comparable to trajectories in the main task (see Supplementary Note 5 and Supplementary Fig. 2). Analogous to the inconsistency levels measured in the value task, we measured subjects’ imprecision in the non-value tasks by calculating the Euclidean distance between the predefined target coordinates and the coordinates of the actual location of the button press (Fig. 2c). Crucially, even though the imprecisions in the non-value tasks were rather small, they captured simple motor errors in task execution. We therefore correlated the Euclidean distances with inconsistency levels in the risky-choice task. We found that in the numerical task, Euclidean distances positively correlated with inconsistency scores in the risky-choice task, suggesting that a greater imprecision in that task was tied to higher inconsistency levels in choice (Behavioral study: β=0.0005, p=0.0063; Replication study: β=0.0008, p<0.0001, OLS regression with random intercepts for subjects). In contrast, in the motor task we obtained a similar result only in the replication study (Behavioral study: β=-0.0002, p=0.567; Neuroimaging study: β=0.0012, p=0.2466; Replication study: β=0.0017, p<0.0001. OLS regression with random intercept for subjects). This shows that motor imprecisions that were measured at the end of motor-only trials were only somewhat related to choice inconsistencies in the risky-choice task. Hence, to fully tackle how motor output from the non-value tasks was related to inconsistency levels in the risky-choice task, we next examined the motor dynamics recorded in these tasks.

As in the main value-based risky-choice task, we repeated the dimensionality reduction procedure using an elastic net regularization on the mouse features extracted from the non-value tasks and tested whether these features correlated with inconsistency levels estimated in the value-based risk task (Fig. 4b and Methods). Since mouse features in the non-value tasks did not involve any value calculations, they can be regarded as strong out-of-sample predictors of the effect that motor dynamics had on inconsistency levels (Fig. 4b). Importantly, we found that the selected mouse features from the non-value tasks accounted for variation in inconsistency levels in the main risky-choice task. That is, mouse features from the motor task explained 3.9%, 6.0%, and 9.9% of the variance in inconsistency scores (behavioral, neuroimaging, and replication studies, respectively; model with random intercepts by subjects: 18.6%, 23.4% and 29.4%), whereas mouse features from the numerical task explained 4.5% and 6.6% of the variance (behavioral and replication studies; model with random intercepts by subjects: 19.0% and 27.2%) (Fig. 4d). Note, that even though the adj-R^2^s are not as high as when using the mouse features extracted from the main risky-choice task, they are still surprisingly high, given that the regressors are derived from a different task.

Our unique analysis approach, which leverages the mouse features, enabled us to point at the exact motor elements that influenced inconsistency scores. Namely, we found that only one feature, the maximal distance from the straight line between the axes-origin and choice (*MD*), significantly correlated with inconsistency levels across the three studies in seven out of the eight different tasks (3 tasks in the behavioral of replication studies and 2 in the neuroimaging study). In all analyses, we obtained a positive coefficient for this feature, suggesting that the larger the distance, the higher the inconsistency levels. This suggests that subjects who tended to wander around the grid further from the shortest path to their choice, regardless of which task they performed, were more likely to be inconsistent. This finding resonates classic findings from motor cognition, which indicate that spatial deviations from straight paths lead to greater motor errors^49^. See Fig 3a for a visualization of the MD feature.

Furthermore, we also found that the average velocity during the trial (*meanVel*) negatively correlated with inconsistency levels in six out of the eight tasks, implying that fast movements led to consistent choices, perhaps indicating that the subjects were precisely aiming at a predetermined specific location. This finding might seem rather contradictory to the canonical speed-accuracy trade-off^23^, where one would expect higher error rates with faster movements. Nevertheless, Decision Field Theory offers a structural explanation to our findings: once discriminability between choice options is low, as in the case in the continuous budget set – an inverse relationship is to be expected, thus faster movements would result in fewer errors^50^. See Fig 3c for visualization of the meanVel feature.

Next, to obtain a true cross-tasks prediction, we treated the regression of mouse features from the main risky-choice task on inconsistency levels as a training set (the same regression model reported in the previous section). We used the estimated coefficients of the mouse features from this regression (*β_k_*) and applied them to be the coefficients of the mouse features extracted from the non-value tasks, which we considered as our test set. Based on this, we calculated the predicted inconsistency levels for the main risky-choice task using the mouse features extracted from either the motor or numerical tasks (*Ŷ* or *Ẑ*, for the motor and numerical tasks, respectively, Fig. 5a). We then correlated these predicted inconsistency levels with the actual MMI scores and found a significant positive correlation in all studies (behavioral study: r=0.462, CI=[0.437;0.485], and r=0.487, CI=[0.463; 0.509], p<0.0001, motor and numerical tasks, respectively, Fig. 5b. Neuroimaging study: r=0.467, CI=[4.38; 4.95], p<0.0001, motor task, Fig. 5c. Replication study: r=0.5895, CI=[0.566;0.612], and r=0.574, CI=[0.550; 0.597], p<0.0001, motor and numerical tasks, respectively, Fig. 5d). This strengthens our findings that specific mouse features from the non-value tasks could predict inconsistency levels in the risky-choice task even when using regression coefficients that were trained on a different dataset.

**Fig 5.**
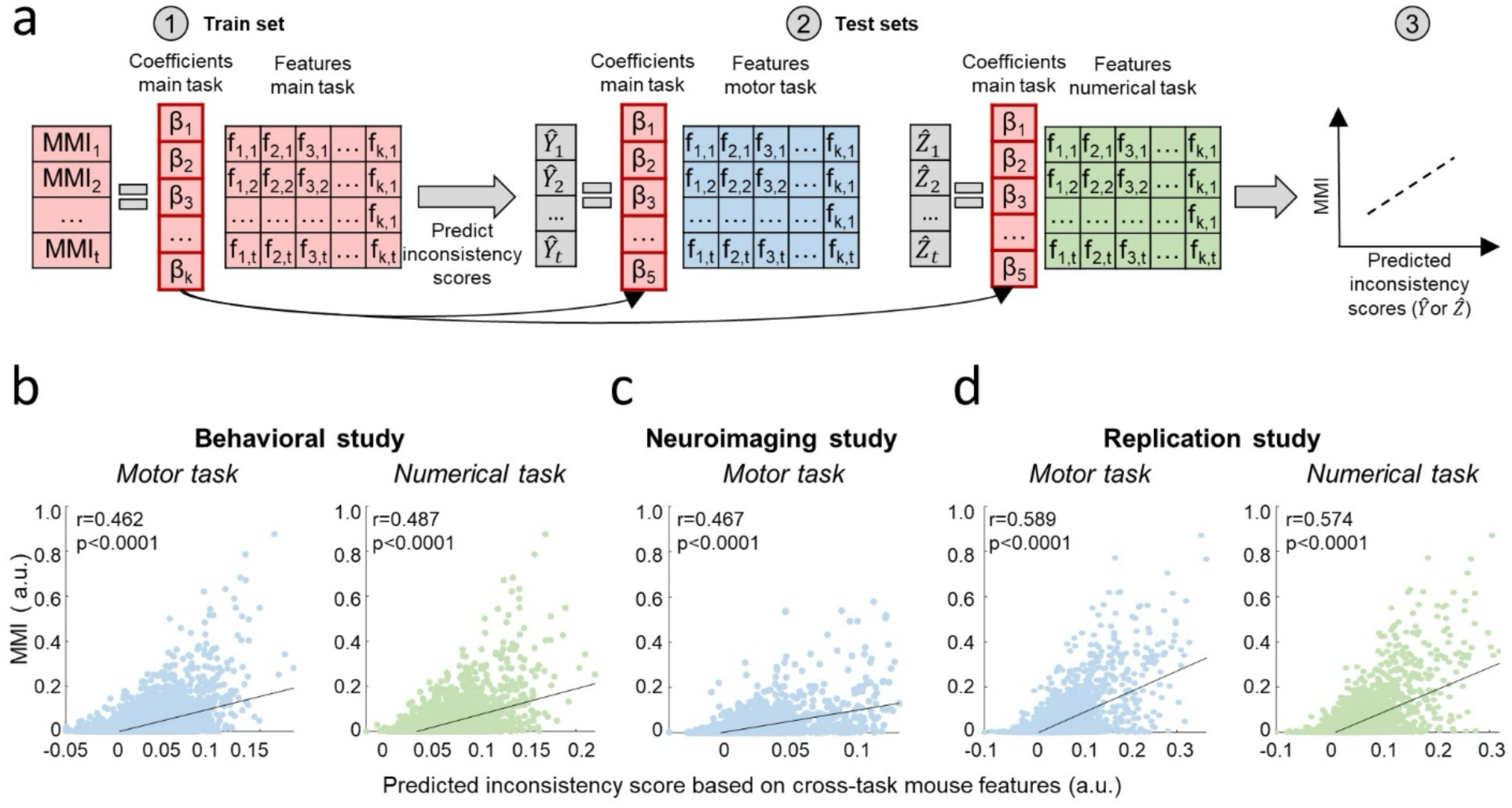
Cross-tasks prediction. (a) We use the coefficients from a regression of MMI scores on mouse features from the value-based task to predict inconsistency scores with mouse features from the non-value tasks. We then correlate predicted scored with actual MMI scores. (b-c) Obtained correlations between predicted and actual MMI. Each dot in the scatter relates to a different trial. (b) Behavioral study. *Left:* predictions based on mouse features from the motor task (N=4,279). *Right:* predictions based on mouse features from the numerical task (N=4,192). (c) Neuroimaging study (N=2,915). (d) Replication study. *Left:* predictions based on mouse features from the motor task (*N=3,085*). *Right:* predictions based on mouse features from the numerical task (*N=3,160*). See supplementary Fig. 5c for the results for *Approach B: PCA*.

Finally, here, too, we repeated the PCA analysis for all the results reported in Fig. 4d-e and Fig. 5 and found similar findings (Supplementary Note 5 and Supplementary Fig. 5b, d).

Taken together, using two different tasks that captured motor dynamics but did not involve value modulation, in three different samples with two different analysis approaches including two out-of-sample predictions, we were able to show that choice inconsistency in value-based decision-making was affected by motor dynamics and computations. Our findings suggest that specific elements in motor dynamics, unrelated to value computation, contributed ∼7% (on average) to the variability in inconsistency scores.

### Neuroimaging results

#### Replication of the main finding from our previous study

As we hypothesized that neural noise in value brain regions during valuation leading up to a choice is one source for choice inconsistency, we first aimed to replicate the main finding from our previous study^27^, which showed that value-modulation and inconsistency levels correlated with the BOLD signal in the same value-related ROIs. We therefore ran the same random-effect general linear model (RFX GLM) used in that study (see Methods) and focused our investigation on pre-hypothesized value-related ROIs. Indeed, we found that MMI correlated with all predefined value ROIs: the ventromedial prefrontal cortex (vmPFC), the ventral striatum (vStr), the dorsal anterior cingulate cortex (dACC) and the posterior cingulate cortex (PCC) (p<0.05, cluster-size corrected, Fig. 6a). We further found a conjunction between regions that correlated with inconsistency and with the SV of the chosen bundle in each trial in three of these regions (vStr, dACC and PCC, p<0.05, cluster-size corrected, Fig. 6a). We did not find similar activations in the whole-brain analysis, perhaps since eight of the subjects in the new sample (19.5%) were fully consistent, which resulted in a weaker correlation of inconsistency levels with the BOLD signal. Another possible explanation is the fact that the current study had lower power^42^ compared to the previous study, as each subject completed only 75 trials instead of 108. Therefore, we also ran a less stringent model, which only included reaction time and inconsistency levels as regressors (Methods). Here, we found a positive correlation between MMI and the BOLD signal in the ACC/mPFC in a very similar region to the one we found in the whole-brain analysis in our previous study (p<0.005, cluster-size corrected with 1,000 Monte Carlo iterations, Fig. 6d). These results indicate that we were able to successfully replicate our previous findings in a different sample, confirming that inconsistency emerges during value modulation.

**Fig 6.**
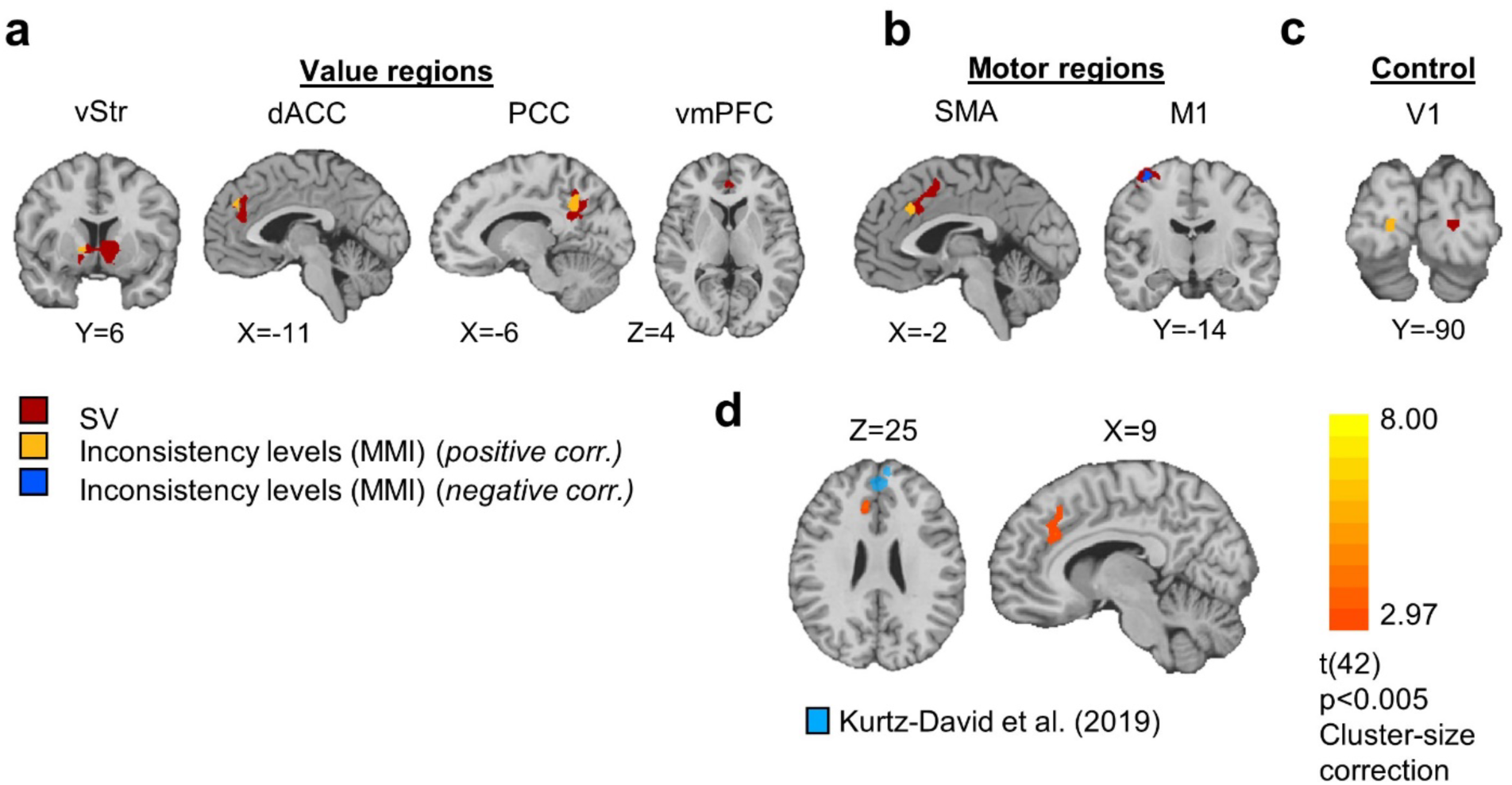
Choice inconsistency correlates with value and motor neural activations. (a-d) An overlay of the MMI and SV regressors. (ROI RFX GLM, n = 42, p<0.05, cluster-size correction. Model regression: *BOLD* = *β*_0_ +*β*_1_*RT* + *β*_2_*MMI_trial_specific_* + *β*_3_*SV* + *β*_4_*Slope + β*_5_*endowment + ε*. Six additional motion-correction regressors were included as regressors of no interest. (a) Replication of the main finding in Kurtz-David et al. (2019)^27^ in value-related ROIs: vStr, dACC, PCC, vmPFC. (b) motor-related brain regions: M1 and SMA. (c) Control ROI: v1. (d) A whole-brain analysis reveals inconsistency levels correlated with the same region found in Kurtz-David et al. (2019)^27^ (blue). Model regression: *BOLD* = *β*_0_ +*β*_1_*RT* + *β*_2_*MMI_trial_specific_* + ε. Six additional motion-correction regressors were included as regressors of no interest. N = 42, p<0.005, cluster-size correction with 1,000 MC iterations. Results are shown on the Colin 152-MNI brain.

#### Motor brain regions also correlate with value modulation and inconsistency levels

Next, we looked at the second hypothesized source of choice inconsistency: neural noise originating from motor planning and execution. We thus tested whether inconsistency levels and valuations correlated with activations in motor brain regions. We ran the same RFX GLM that we conducted in the value-related brain regions, this time on M1 and the supplementary motor area, SMA (see Methods and Supplementary Fig. 6), and found that both these regions correlated with inconsistency levels (SMA – positive correlation, M1 – negative correlation, p<0.05, cluster-size correction, Fig. 6b). In line with previous studies, we also identified a strong activation of the SV regressor in these regions, which demonstrates the notion that motor regions receive information regarding valuations from value-related brain areas before execution^51^, and/or that this activation is evidence for the representation of action values^52^.

Here, too, we saw an overlap between the neural correlates of SV and inconsistency levels (p<0.05, cluster-size corrected, Fig. 6b), suggesting that not only did activity in motor brain regions encode valuations of choice, but rather also tracked, and perhaps contributed to inconsistency levels.

Finally, we also examined the activity in V1 as a control region and found a very small cluster in the right hemisphere, which correlated with MMI, and another very small cluster in the left hemisphere, which positively correlated with SV. Nonetheless, given the contralateral correlates of the two regressors, there was no conjunction between them, nor did these findings survive a more stringent threshold (see Methods and Fig. 6c). This suggests that valuations and inconsistency levels were not jointly represented in any main brain region to be tested.

#### Functional connectivity between motor and value brain regions moderates inconsistency levels

If, in fact, both value and motor brain regions contribute to choice inconsistency, then inter-transmission activations between these regions should trace inconsistency levels. Therefore, we performed a psychophysiological interaction (PPI) functional connectivity analysis (see Methods). We used the value regions (vmPFC, vStr, dACC, and PCC) as seeds and ran a separate PPI regression for each of the two motor regions (M1 and SMA; for a total of 8 models). We found that as MMI increased (more inconsistent), M1 had stronger connectivity with three of the value regions (vStr, dACC and PCC; see Supplementary Table 6). In the SMA, we found a similar result for the dACC seed. These results show that the activity in motor and value brain regions were more synchronized in inconsistent trials than in consistent trials, strengthening the notion that motor-related computations also contributed to inconsistency levels.

#### Neural correlates of mouse features

We next examined how the mouse features were represented in the brain. Our main goal was to compare the neural correlates of the mouse features during the value-based risky-choice task to the non-value motor task, and then test whether this representation was related to the degree of inconsistency levels. We first examined the neural correlates of mouse features in both value and motor ROIs. We hypothesized that the mouse features would correlate with activity in the motor ROIs in both tasks, as these features reflected the motor dynamics of task execution regardless of the specifics of the task at hand. However, we speculated that mouse features would be evident in value-related ROIs solely during the main value-based risky-choice task, where they also reflected the valuation process.

To test this, we ran a RFX GLM on the BOLD signal in the value and motor ROIs using the three mouse features, which significantly correlated with MMI in both the main and motor tasks of the neuroimaging study: mean velocity during the trial (meanVel), the maximal distance from the shortest path to the choice (MD), and the number of segments in which the subject ceased motion during the trial (numFixations) (Fig. 4c middle panel, and Supplementary Table 4). Note that contrary to MD and meanVel, which were strong predictors of inconsistency levels across all three studies (see above), numFixations was a strong predictor of inconsistency levels solely in the neuroimaging study (in both tasks). Fig. 7a presents an overlay of the neural correlates of the mouse features in all ROIs in the risky-choice task (Fig. 7a, upper row) and motor task (Fig. 7a, bottom row).

**Fig 7.**
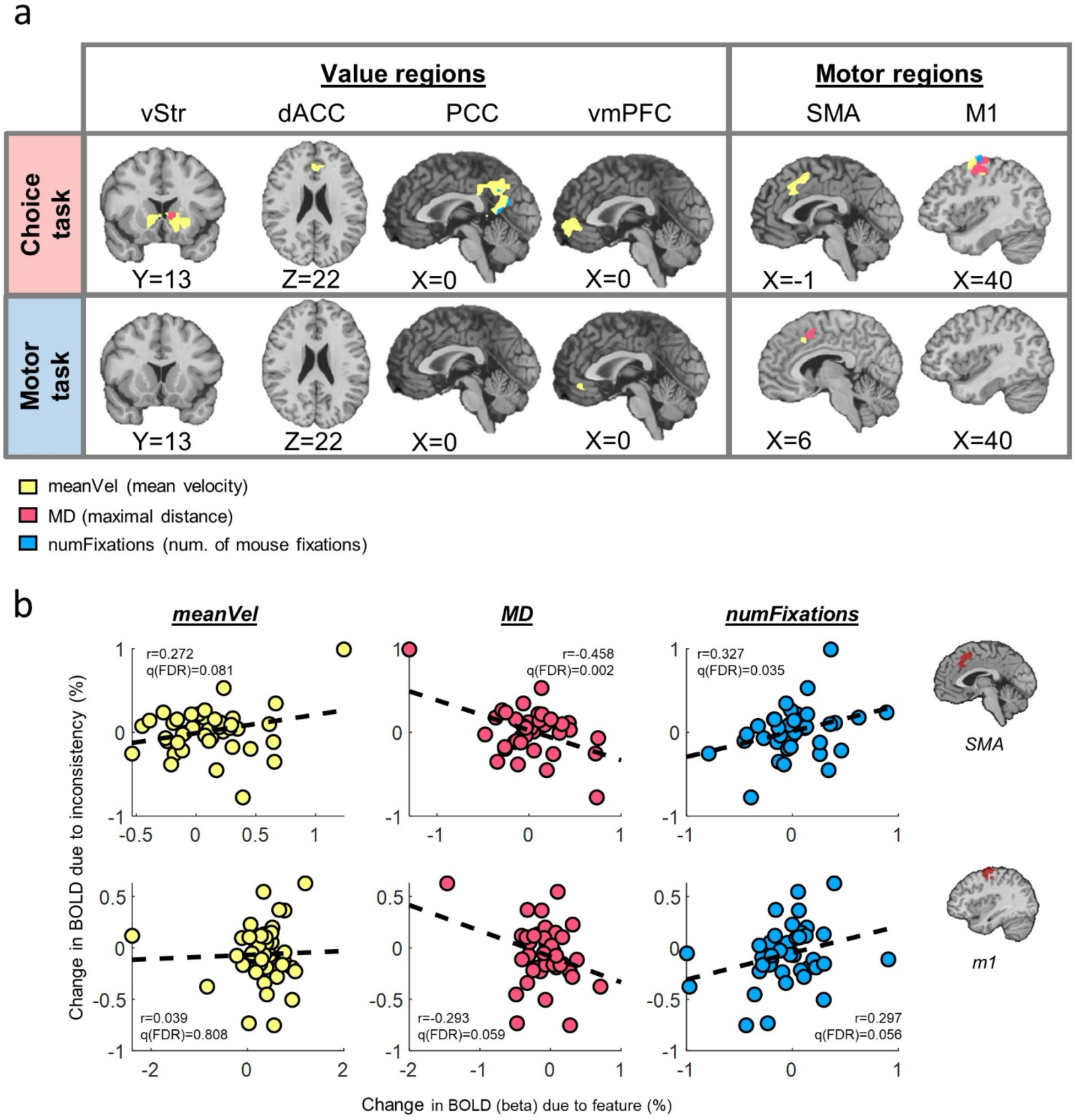
Neural correlates of motor dynamics (*elastic net based features)*. (a) An overlay of the neural correlates of motor features, which were selected by the elastic net analysis. *Upper row:* main risky-choice task. *Bottom row:* motor task. Left column presents value-related ROIs, whereas the right column presents motor ROIs. ROI RFX GLM, n = 42, p<0.05, cluster-size correction. Model regression: *BOLD* = *β*_0_ +*β*_1_*RT* + *β*_2_*meanVel* + *β*_3_*MD* + *β*_4_*numFixations + β*_5_*slope + β*_6_*endowmnt + ε*. Results are shown on the Colin 152-MNI brain. (b) Neural activations of mouse features in motor brain regions correlated with the neural representation of inconsistency levels. We obtained the beta coefficients attributed to each mouse feature from a trial-by-trial RFX GLM (representing the change in BOLD signal, model presented in Fig. 7a, upper row), as well as the beta coefficients attributed to inconsistency levels from the RFX GLM of the BOLD signal on MMI (model presented in Fig. 6b). We then correlated the two sets of coefficients on the subject-level. Each scatter represents subject’s random effects (slopes) from the two different models, and refers to the change in BOLD activation associated with mouse features (x-axis) compared with the change associated with inconsistency scores (y-axis). Up: SMA ROI, bottom: m1 ROI.

Not surprisingly, we found that mouse features correlated with activations in the motor ROIs in both the main task and the motor task (p<0.05, cluster-size corrected). However, in the value ROIs we found a large discrepancy between the two tasks. While in the risky-choice task we identified that the mouse features strongly correlated with activations in value ROIs (clusters of activations that correlated with mouse features in each ROI: vStr: 3/3 features, dACC: 1/3 features, PCC: 2/3 features, vmPFC: 1/3 features, p<0.05, cluster-size corrected), we barley found any representation of the mouse features in value ROIs during the motor task (clusters of activations that correlated with mouse features in each ROI: vStr: 0/3, dACC: 0/3, PCC: 0/3, vmPFC: 1/3, p<0.05, cluster-size corrected). In other words, in line with our hypothesis, neural activations associated with motor dynamics during the motor task were much weaker in the value ROIs than during the risky-choice task. Hence, only during the risky-choice task, we found that choice dynamics similarly activated motor and value brain regions. This suggests that there was a transmission of motor cues between the two networks, only when task-execution involved a valuation process. In contrast, given that all three features were shown as robust predictors of inconsistency levels in the non-value tasks, the activations associated with these features in the motor brain regions during the motor task is another indication that motor-only activations contributed to inconsistency.

We next aimed to directly relate the neural activations of choice dynamics in the motor brain regions with inconsistency levels, to examine if the relationship we found in behavior between choice dynamics and inconsistency levels also holds at the neural level. To this end, we obtained the beta coefficients from the RFX GLM that modeled the trial-by-trial change in the BOLD signal in M1 and SMA that was attributed to each mouse feature (subjects’ random slopes, risky-choice task, presented in Fig. 7a, upper row), as well as the beta coefficients from the RFX GLM that modeled the trial-by-trial change in the BOLD signal attributed to inconsistency levels (MMI) in the same ROIs (subjects’ random slopes from the model presented in Fig. 6b). We then correlated, on the subject-level, these two sets of beta coefficients. A significant correlation between the two sets of beta coefficients would suggest that activations associated with inconsistency levels were also associated with a specific aspect of the motor movement captured by the mouse feature under examination.

We found significant correlations for two of the three mouse features we examined (MD and numFixations) in SMA, and marginally significant correlations for the same two features in M1 (MD: SMA: r=-0.458, q(FDR)=0.002, CI=[-0.669;-0.179]; M1: r=-0.293, q(FDR)=0.059, CI=[-0.548;-0.0114]. numFixations: SMA: r=0.327, q(FDR)=0.035, CI=[0.025;0.574]; M1: r=0.297, q(FDR)=0.056, CI=[0.010;0.622], Fig. 7b). The smaller effect-size in M1 might be related to the weaker inconsistency signal in that ROI (Fig. 6b).

To conclude, the current analysis shows that motor characteristics of task execution had an explicit neural footprint in motor circuits, one which directly affected the neural computation of inconsistent choice behavior. This provides further evidence for the role of the motor cortex in inconsistent choice in value-based decision-making.

#### Ruling out value modulation in the motor task

Finally, to validate that our non-value motor task did not involve any sort of value modulation related to choice, we conducted two separate analyses. First, we conducted a pseudo-SV analysis, where we treated the location of the button-press in each trial as if it was an actual choice had it been conducted in the value-based risky-choice task. Based on this assumption, we calculated what would have been the inconsistency level and SV had it been a choice in the value-based task (Methods and Fig. 8a). We then looked for the neural correlates of this pseudo-SV regressor with the BOLD signal in the motor task, using the same RFX GLM modelling that we used in the analysis described in Fig. 6. We found null results for all predefined value ROIs (Fig. 8b, p<0.05, cluster-size corrected), indicating that subjects in the motor task did not conduct monetary-based value computations during motor execution, nor did they treat the location of the button-press as choosing a lottery.

**Fig 8.**
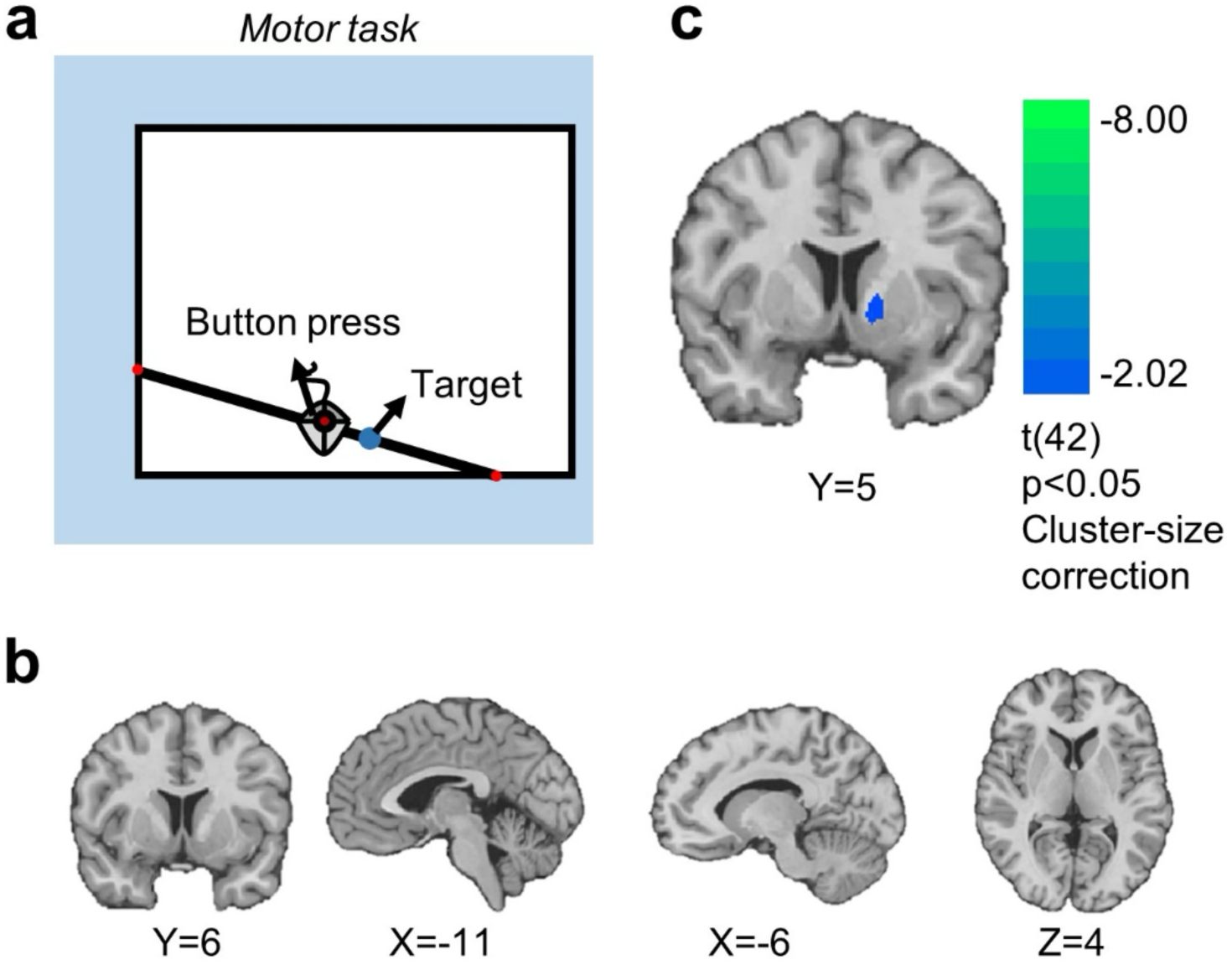
Ruling out value modulation in the motor task. (a-b) pseudo-SV analysis. (a) We treated the button-press location in the motor task as if it was a choice in the value-based risky-choice task. Based on this assumption, we then extracted utility-parameters, corresponding to this location, and calculated what would have been its trial-specific-MMI score and trial-specific-SV had it been a lottery choice in the value-based risky-choice task. (b) We repeated the same ROI RFX GLM from Fig. 6a-c on the BOLD signal from the motor task with the pseudo-MMI and pseudo-SV regressors. (c) A reinforcement signal for reaching the target in the motor task. For each TR, we calculated the current Euclidean distance from the target. We then ran a RFX GLM in value-related regions. n=42, p<0.05, cluster-size correction. Model regression: *BOLD* = *β*_0_ + *β*_1_*RT* + *β*_2_*EucDistance* + *β*_3_*slope* + *β*_4_*Y_Intersection + ε*. Six additional motion-correction regressors were included as regressors of no interest. Results are shown on the Colin 152-MNI brain.

In the second analysis, we hypothesized that the motor task carried a different type of decision-making procedure, one that did not include value computations, but rather a different type of computation. We speculated that at each point in time during the trial the subject determined the distance between the mouse curser and the predefined target, indicating whether they had already reached the predefined target’s coordinate (Methods). To this end, in each repetition time epoch (TR), we calculated the Euclidean distance between the current position of the mouse’s curser and the predefined target and looked for its neural correlates in all value-related ROIs. We found it was negatively correlated with BOLD activations in the vStr, meaning, that the vStr became more active as subjects were closer to the target, probably suggesting a reinforcement-like signal for target-reaching (Fig. 8c, p<0.05, cluster-size corrected). This result shows that during the motor task subjects did not calculate subjective value nor tried to maximize utility of chosen lotteries, but instead calculated the spatial location of the mouse relative to the target location.

## Discussion

In this study, we used behavioral paradigms, neuroimaging, and mouse tracking techniques to show that suboptimal choice is the result of two interacted neural processes. Idiosyncratic choices, leading to violations of fundamental economic axioms, rise during the noisy process of value computation, but at the same time are also influenced by mechanisms of task-execution instantiated by motor output. The current research applied mouse-tracking techniques to a well-established task in risky choice, and presented innovative non-value motor tasks that isolated motor planning and execution from choice dynamics in the same experimental setting. The findings reported in the current paper were obtained in two very different settings (computer vs inside the MRI scanner), and then replicated in a third separate sample.

Similarly to previous studies using the same task^27,28,32^, we found heterogeneity in subjects’ inconsistency levels, and that subjects did not choose at random (Fig. 2a-b). In the family of random utility models^53–55^, choice inconsistency is the result of random fluctuations of the utility function. However, importantly, this embedded stochasticity does not lead to randomness in choice – i.e., any value could be drawn at any time with an equal probability. We directly demonstrated this notion by comparing our subjects to simulated random decision-makers (Fig. 2a). Thus, for the most part, choosers obeyed monotonicity and transitivity, but fluctuations in neural computations led, in some cases, to inconsistent choices. We therefore examined two sources for such fluctuations: the valuation process per se, as well as the motor dynamics of choices.

Through our novel task design, we were able to show that motor imprecisions in the non-value tasks were partially related to inconsistency levels in the main risky-choice task. However, to provide a well-rounded report of the role of motor output in inconsistent choice, a far more thorough investigation of motor dynamics was required. Hence, we extracted multiple (34) mouse features from recorded trajectories to characterize temporal and spatial variation in motor dynamics. Thus, we did not focus solely on a small set of features that had been previously shown to be related with valuation and choice (e.g. mean velocity^38^ or the curvature of the trajectory^39,56^). Controlling for other task parameters, the mouse features from the risky-choice task accounted for roughly 7-18% of the variation in inconsistency levels. Crucially though, we obtained a close estimate (∼7% on average) when using mouse features that solely originated from the non-value tasks. As these tasks did not involve any value-related computations (Fig. 8), they solely represented motor planning and execution. We illustrated this finding in two different analysis approaches, with both in- and out-of-sample prediction techniques (Figs. 4-5).

The contribution of mouse features to inconsistency levels was larger in the behavioral and replication studies (>16%) than in the neuroimaging study (7%), probably because subjects’ movement was not restricted, and they could freely move during the study. This resulted in a greater role of volatile motor computations in inconsistent choice. Along these lines, previous studies claimed that the contribution of motor noise to variability in choice was much more limited (3-5%)^1,11^. We believe that this discrepancy is rooted in the far more complex motor output in our task – moving the mouse towards a budget line vs. a button-click in 2-alternative forced choice paradigms in previous studies. We pose that such designs were too simple to fully examine the complexities of the motor system and rigorously test if motor noise contributed to choice inconsistency. Hence, we consider the complex motor dynamics in our task as a strong advantage, which allowed us to thoroughly tackle the role of motor noise in choice inconsistency.

The only feature that correlated with inconsistency levels across all tasks and samples was MD, the maximal distance from the shortest path to the final choice, which indicated that subjects, who travelled in straight “decisive” pathways towards choice, were less likely to demonstrate high inconsistency levels. This suggests that these routes had lower noise levels and corroborates the groundbreaking work of Harris & Wolpert (1998)^20^. They argued that straight trajectories are selected to minimize the variance in end-point positions, as a result of a unimodal velocity profile that minimizes the accumulated signal-dependent noise, which is the cause for deviations from targets^20^. In our study, minimizing the variance towards the end-point also resulted in lower inconsistency levels. Since the intensity of inconsistency levels is sometimes referred to as a measure of wasted income^41^, these routes can be considered less costly, suggesting that motor planning takes into account subjects’ own motor uncertainty which carries monetary costs^57,58^.

We were able to replicate our previous neuroimaging results^27^ and show that inconsistency emerged during the computation of value and was represented in value related brain areas, but at the same time, was also affected by activations in motor brain regions (Figs. 6-7). Our neuroimaging results further tied between neural activations associated with specific elements in task execution to inconsistent choice. This implies that the decision output in our task was not only the result of value computations in value brain regions, but was also affected by the motor system. Several studies have already shown how noisy neural computations of value, rather than explore-exploit or diversification strategies, may cause irrationality or imprecision in value-based choice^11,27,59–61^. Similarly, spontaneous neural activity also accounted for stochastic behavior in sensory and motor decision tasks that did not involve valuation processes^1,14,62^. Here, we associated motor activations directly to value-based choice, and argued that the neural dynamics that give rise to inconsistent choice were not limited to the value system and go beyond suboptimal valuations. Our results highlight the importance of studying the role of motor noise outside classic paradigms of motor skills (i.e. in grabbing or target reaching).

Our neuroimaging findings further indicated that choice dynamics correlated with activations in value ROIs during the main risky-choice task, suggesting that mouse features reflected fundamental aspects of the valuation process by itself, ones that went beyond motor planning and task execution. The weaker correlations during the motor task further indicated that the bidirectional transmission between the networks was less necessary in the absence of a value-based choice process. This can imply that the transmission between decision circuits and motor regions was perhaps not sequential nor was it only feed-forward. Rather, it may reflect a continuous flow of information between circuits^63^, which can point at further deliberation after movement onset^64^.

We study decision mechanisms and task execution using mouse tracking, which captures temporal and spatial dynamics with high resolution. Choice dynamics also illuminate deliberation processes between chosen and unchosen options, while indicating the timing and loci of subtask computations^36^. Thus, mouse tracking has been extensively used in recent years to study implicit cognitive mechanisms in several domains, such as social categorization and face recognition^65–67^, psycholinguistics^68,69^, visual search^70^, and memory recognition^71^. Lately, mouse tracking has also been applied to the study of value-based decision-making, including consumer goods and dietary choices^38–40^, learning mechanisms^72^ and inter-temporal choice^56,73^. This line of studies was followed by rich and exhaustive methodological reports^36,37,43–46,74,75^, but, to date, has focused only on binary choice tasks. Our study builds upon the standard mouse-tracking methodological framework and expands it to a task with a continuum of multiple-choice options. Our metrics can be applied to many other tasks, which use a wide array of choice options.

Economists have long argued that inconsistencies in subjects’ responses can be the result of some random components in their preferences ordering^76,77^. This led to calls for improved theories of how stochastic noise influences behavior^78,79^. More recently, these inconsistencies were modeled as resulting from a deterministic source^80^. The “constant-error” approach^81^ suggested that subjects obeyed some deterministic theory of choice, but would experience a tremble or a lapse with some probability at the execution of choice. In contrast, the “white noise” approach posed that subjects maximized a utility function with some additive noise term^55,82^, implying that stochasticity in choice occurred during the valuation process. This approach was supported by neural evidence, suggesting that the source for these stochastic valuations was the spontaneous fluctuations of neural networks in value-related brain regions^17,27,61^. Our findings can help refine theory, and enable a finer description of the magnitude and form of the noise component^82,83^. Lately, Webb^17^ has shown that “white noise” models are analogous to sequential sampling models, often used in psychology and neuroscience to describe dynamic choice processes in discrete – usually binary – choice sets. The current study presented vast empirical evidence that bridges these schools of thought and suggests that irrationality can be the result of a combination of two different random neural processes.

Finally, one can easily think of many real-world decisions for which our findings may be relevant. For example, decisions during driving require constant updating of motor planning and execution circuits. Likewise, mobile or web e-commerce, as well as decisions that make use of sliding scales, such as for donations or liking ratings – all involve substantial motor element within task execution. Going back to the arcade analogy, we show that even picking up your favorite teddy bear out of the claw machine, may be diverted with imprecise motor implementation.

## Methods

### Stimuli

#### Main risky-choice task

Subjects made choices from linear budget sets in the context of risk^27,28^. On each trial, subjects were presented with a set of 50/50 lotteries between two accounts, X and Y, and were asked to choose their preferred lottery (a bundle of some amount of X and some amount of Y). All possible lotteries in a given trial were represented along a budget line. The price ratios (slopes of the budget lines) and endowments (distance of the budget line from the origin) were randomized across trials (see Supplementary Tables 1 and 2 for budget sets’ parameters). Importantly, we set the mouse cursor at the axis-origin at the beginning of each trial, and recorded mouse trajectories (see Mouse Tracking below). For further details on the main task, see the left panel of Fig. 1a and our previous study^27^.

#### Non-value motor tasks

Following the main task, subjects conducted non-value motor tasks, in order to examine motor-related sources of choice inconsistency that did not involve value modulation. We used these tasks to test whether certain characteristics in subjects’ motor planning and execution could account for subjects’ choice behavior and inconsistencies.

##### Motor task

Subjects were presented with linear graphs and had to reach a black circular target on the graph, resembling target-reaching tasks. The graphs, nor the axes-grid, presented numbers. When subjects reached the black target, they left-clicked the mouse to submit their position (middle panel of Fig. 1a).

##### Numerical task

Similarly, the numerical task tested subjects’ motion pathways when absent value modulation, though instead of visual circular targets, here, subjects had numerical coordinates as targets, resembling spatial navigation tasks. Subjects were presented with linear graphs, and had to reach a predefined {*x*, *y*} coordinate appearing at the top of the screen, which represented a target spot on the graph (right panel of Fig. 1a). The current {*x*, *y*} cursor position was presented continuously at the top of the screen. When subjects reached the target coordinate, they left-clicked the mouse/trackball to submit their position.

In both non-value motor tasks, to fully resemble the main task, each graph (both the slope and the endowment) was identical to a graph from the main risky-choice task. The predefined targets for both non-value motor tasks were identical to the actual location on that graph that each subject chose in the main risky-choice task. That is, the loci of the predefined targets were subject-specific. Note that in both tasks there was no value-based decision-making, as the X- and Y-axes did not represent monetary payoffs. Here, too, we set the mouse cursor to the axes-origin at the beginning of each trial, and recorded mouse trajectories. Supplementary Note 1 provide English versions of the instructions sheet of the non-value motor tasks (behavioral study).

#### Experimental procedures Behavioral and replication studies

*Participants.* Behavioral study: we recruited 91 volunteering undergraduate students from various departments at Tel Aviv University (46 females, mean age 24.2, 18-28). Our sample-size was determined based on previous studies using the same task^28,29,32^. In the main task, two subjects were dropped from the sample due to technical problems during their run. We report the results for the remaining 89 subjects. Five subjects in the numerical task and six subjects in the motor task were dropped due to technical problems in their computer stands during the run. We report the results from the remaining 85 and 86 subjects, respectively.

##### Replication study

The desired sample-size was determined based on a power analysis that used the *r* correlation coefficients from cross-tasks prediction presented in Fig. 5B, which yielded an ES of r=0.462 and r=0.487 (motor and numerical tasks, respectively). We computed the desired-sample size using the smaller ES (r=0.462), implemented in the standard G*Power software (correlation: bivariate normal model, one-sided). We concluded that to achieve a power of 90% with α=5%, we had to recruit at least 37 subjects. In practice, we recruited 72 volunteering undergraduate students from various departments at Tel Aviv University. In the main task, three subjects were dropped from the sample due to technical problems during their run. We report the results for the remaining 69 subjects (49 females, mean age=28.9, 18-60). Three additional subjects in the numerical task and seven subjects in the motor task were dropped due to technical problems in their computer stands during the run. We report the results from the remaining 66 and 62 subjects, respectively. All the sessions of the behavioral and replication studies were carried out in the Interactive Decision-Making Lab. All subjects gave informed written consent before participating in the study, which was approved by the local ethics committee at Tel-Aviv University.

##### Sessions

*Main task.* Before the main task, subjects read an instruction sheet for that task. The instructions were repeated aloud by the experimenter. Subjects then carried out a short practice on the experimental software, and went on to the main task. Subjects made a total of 150 trials, divided into three blocks of 50 trials each (see budget sets’ parameters in Supplementary Table 1). On each trial, subjects had a maximum of 12 seconds to make their choices, followed by a 1 sec variable inter-trial-interval (ITI, jittered between trials).

##### Non-value tasks

Following the main task, we randomly chose 51 budget sets (out of 150) to be used as the graphs in the two non-value tasks. Subjects then received the instructions sheet for the two additional tasks, and went on to complete the two tasks. The order of the tasks was counter-balanced between-subjects across experimental sessions. Within-subjects, trials were presented at a random order within each non-value task.

In the motor task, subjects had up to 6 sec on each trial to reach the predefined black target, followed by a variable ITI of 1 sec (jittered between trials). In the numerical task, subjects had a maximum of 12 sec on each trial to reach the predefined {*x*, *y*}coordinates, followed by a variable ITI of 1 sec (jittered between trials) (see Fig. 1b, upper panel).

##### Payoffs

At the end of the experiment, one of the trials from the main risky-choice task was randomly selected for monetary payment. Subjects tossed a fair coin to determine the winning account, *X* or *Y.* Subjects won the monetary value of the tokens allocated to the winning account on the trial drawn for payment. Each token was worth 1 NIS (at the time, $1 ≅3.5 NIS).

At the end of each non-value motor task, we casted one trial at random, and compared the predefined target coordinate with subjects’ actual submitted position. Subjects received 5 NIS for each non-value motor task, though this amount decreased as a function of their Euclidean distance from the target. We implemented this payment method to incentivize subjects’ precision and motivate them to reach the predefined targets. By doing so, we could ensure that trajectories from the non-value motor tasks would be comparable with those in the main task (Fig. 1e). The average prize was 40.5 NIS (behavioral study) and 42.7 NIS (replication study). Subjects also received 25 NIS as a show-up fee.

#### Neuroimaging study

##### Participants

We recruited 46 right-handed volunteering students from various departments at Tel Aviv University (22 women, mean age 25.5, 19-49). Sample-size was determined based on our previous study^27^. Subjects gave informed written consent before participating in the study, which was approved by the local ethics committee at Tel-Aviv University and by the Helsinki Committee of Sheba Medical Center. We dropped scans with sharp head movements (>3 mm in translation / 3^0^ in rotation). As a result, three subjects were dropped from the analysis of the main task and two subjects were dropped from the analysis of the motor task. For three additional subjects we had to drop one scan (either from the main or the motor task), and for one subject we dropped two scans (one from each task). Another subject was dropped due to technical problems during their session. We therefore report the data for the remaining 42 subjects in the main task and 43 subjects in the motor task.

##### Instructions and pre-scan practice

Before the scan, subjects read an instruction sheet for both the risky-choice and motor tasks (note that subjects did not conduct the numerical task in the neuroimaging study), and thereafter completed a pre-scan questionnaire to verify that the task is clear. For details on the pre-scan questionnaire see Kurtz-David et al. (2019)^27^. Thereafter, subjects completed a practice block of the two tasks in front of a computer, using a similar trackball to the one used inside the fMRI, in order to imitate the motor movements required during the scan. The budget sets in the practice block were predefined to ensure all subjects encountered the same (substantial) variation of slopes and endowments.

##### Sessions

*Main task.* Subjects performed the experimental task using an fMRI compatible trackball to choose their preferred bundle. We let subjects complete another three practice trials inside the MRI scanner to gain experience with the MRI-compatible trackball. Then, the experiment began. Subjects made a total of 75 trials, divided into three blocks of 25 trials each. On each trial, subjects had a maximum of 11 seconds to make their choices, followed by a 6 sec variable ITI (jittered between trials). See below details on image acquisition.

*Motor task.* We used the exact same 75 budget sets and subjects’ choices from the main task as graphs and targets in the motor task (randomized within and across subjects). Subjects had up to 6 sec on each trial to reach the black target, followed by a variable ITI of 4 sec (jittered between trials) (see Fig. 1b, bottom panel). After completing the two tasks, we obtained anatomical scans.

*Payoffs.* The payment protocol was identical to the behavioral study, with small modifications to increase monetary incentives. In the main task, each experimental token was equal to 2 NIS (rather than 1 NIS), and the endowed payoff for the motor task was 20 NIS (rather than 5 NIS). The average prize was 88.6 NIS + 100 NIS show-up fee.

*Image acquisition*. *fMRI.* Scanning was performed at the Strauss Neuroimaging Center at Tel Aviv University, using a 3T Siemens Prisma scanner with a 64-channel Siemens head coil. To measure blood oxygen level-dependent (BOLD) changes in brain activity during the experimental task, a T2*-weighted functional multi-band EPI pulse sequence was used (TR = 1 s; TE = 30 ms; flip angle = 68°; matrix = 106 × 106; field of view (FOV) = 212 mm; slice thickness = 2 mm; band factor = 4). 64 slices with no inter-slice gap were acquired in ascending interleaved order, and aligned 30° to the AC–PC plane to reduce signal dropout in the orbitofrontal area. Each run in the main task had 428 TRs (∼7.1 minutes) and each run in the motor task had 253 TRs (∼4.1 minutes).

*Anatomy.* Anatomical images were acquired using a 1 mm isotropic MPRAGE scan, which was comprised from 176 axial slices without gaps at an orientation of - 30° to the AC–PC plane to reduce signal dropout in the orbitofrontal area.

#### Inconsistency analysis

For a subject-level analysis, we calculated the number of GARP violations^24^. Respectively, we used Afriat Index (AI)^41^ and Money Metric Index (MMI)^32^ to measure the intensity of violations. AI^41^ is a non-parametric index that measures the largest shift to all budget sets (in percentages) required to remove all the violations. It can be interpreted as the subject’s waste of income due to inconsistency. MMI^32^ is a parametric index which measures the minimal shifts to the budget sets (in percentages) required to reconcile incompatibilities between the subject’s choices and its best-fitting parametric utility function, such that all violations are removed from the set. Both indices range between 0 and 1, with higher indices values indicating higher inconsistency levels.

Similarly to our previous study^27^, for a trial-by-trial analysis of inconsistency levels, we implemented a Leave-one-out procedure on MMI to yield trial-specific-MMI. Namely, the index measures the difference between the subject-level MMI (calculated on all the trials in the dataset) and the MMI calculated over all trials except the current trial. See Supplementary Fig. 7, Supplementary Note 3 and Kurtz-David et al. (2019)^27^ for further details and formal definitions.

#### Parametric utility recovery

For each subject, we followed the common-practice in the revealed preference literature^27,28,32,84^ and recovered the subject-specific utility parameters, using Gul’s Disappointment Aversion function^85^. To use subjective value (SV) as a parametric regressor in our behavioral and neuroimaging analyses, we calculated the value of the Disappointment Aversion model with constant relative risk aversion (CRRA) functional form at the chosen bundle (*x^i^*, *y^i^*) in each trial *i*, using the subject’s recovered parameters, *β* and *ρ*, which capture the outweighing of the better option and the subject-specific relative risk-aversion, respectively. See Supplementary Note 3 and Kurtz-David et al. (2019)^27^ for further details on the function. Since the valuation of the chosen bundle might be confounded with motor movement, we used SV as a control variable in the regressions reported in the paper.

#### Measuring choice difficulty

For each subject and trial, we calculated the choice simplicity index (CSI), as presented in our previous study^27^ (See also Supplementary Note 6). Briefly, in each trial, we computed the subjective valuation (SV) of 1,000 lotteries along the budget line, and then computed the variance between these valuations, such that smaller variance indicates a greater difficulty as lotteries along the line are more similar to one another (smaller value difference between the lotteries). Since a high difficulty level of a given trial can be a source for choice inconsistency, we used CSI as a control variable in the regressions reported in the paper.

#### Mouse tracking preprocessing

We recorded mouse movements using MATLAB built-in functions. Each movement of the mouse triggered writing its location in its {*x*, *y*} coordinates and a timestamp in a dedicated excel file. Since sampling occurred only when the mouse moved, it resulted in unequal sampling intervals. We used the timestamps and coordinates to interpolate the mouse location in either 100 or 1,000 samples with equal intervals in time (using MATLAB’s interp1 function). The total time of a trial differed, so the resulting 100/1,000 samples had different sampling rates per trial, but a constant sampling rate within trial. In addition, for some features we required trajectory samples to represent a fixed curve rather than movement through time. Therefore, we also created a version of each trajectory where samples were evenly spaced on the X and Y axes, neglecting the time domain. In this case, the number of samples per trajectory was determined as half the total distance travelled in the trial (in coordinates units), to create a coherent and smooth representation of the trajectory curve. All features reported in this study were extracted on either the evenly spaced trajectory curve, the 100 mouse trajectory samples with equal time-intervals, or the 1,000-samples trajectory, which provided higher temporal resolution and enabled more precise calculations in certain features.

Trajectories with two samples or lower could not be analyzed and were discarded. This was caused by faulty sampling or a hastened miss-click by the subject, and occurred only in 1 trial in the risky-choice task, 13 trials in the numerical cognition task and 29 trials in the motor task (behavioural study). In the neuroimaging and replication studies, this did not occur, so no trials were excluded due to a low number of samples.

#### Extracting mouse features

We characterized each mouse trajectory using multiple features, in order to extract as much information as possible on the decision process and task execution, from various aspects and approaches. Since we did not know ex-ante which features would relate to either value modulation or motor processes, we aimed to extract a large variety of features. Many features were based on or inspired by Jonathan B. Freeman’s extensive work on mouse trajectories, such as maximum deviation (MD), complexity (e.g. Xflips and Yflips), velocity (e.g. meanVel), acceleration (e.g. meanAcc), and area under the curve (AUC), adapted to our experimental design^36,37,43,44,86^. We also calculated the trajectory angle at each time point, with respect to a straight line from the axes origin towards the chosen bundle, adapted from a work by Sullivan and colleagues^38^. Additionally, we used sample entropy^87^ to discern the complexity of trajectories (see Feldman & Crutchfield (1998)^88^ for a discussion of complexity measures). For the purpose of extracting trajectory curvatures at each time point we employed an open-source MATLAB code for curvature calculation, on the evenly spaced trajectory curve^89^. Other features, such as features related to mouse fixations (numFixations) and layover at the axis’ origin (Layover), or features concerned with the time spent in different areas relative to the grid, namely - the time out of the grid bounds (XTimeOutofBounds and YTimeOutofBounds), time above budget line (AboveLine), time near the predicted bundle (TimeNearPredicted) and time on budget line (TimeBudgetLine), were our own design.

We extracted a total of 34 mouse features from recorded trajectories. A comprehensive list of all the features we used appears in Supplementary Table 3, while a more detailed account on the extraction of each feature appears in Supplementary Note 2.

#### Dimensionality reduction

Due to the high number of mouse features relative to the number of trials (both studies), and due to inter-features correlations, we had to induce dimensionality reduction procedures. To this end, we took two different strategies: *Approach A:* feature selection via elastic net, a linear combination of the Ridge and Lasso regularized regression methods; *Approach B:* a feature projection strategy via PCA. The main advantage of the elastic net approach is that it allows for a straightforward interpretation of the results on the expense of losing some information concealed in the features that did not survive the regularization. On the contrary, the PCA approach maintains all mouse-features’ data, but loses some of the interpretability. We implemented these two strategies in mouse features from the main risky-choice task (z-scored) in each of the three studies.

The elastic net analysis was run against inconsistency levels (MMI) and included subjects fixed-effects with 10 folds cross-validation. In the main task, a total of 30 features survived the regularization in the behavioral study (*α* = 0.8, *λ* = 0.0001, with a mean squared prediction error (MPSE) of 0.0034). A respective group of 30 features survived the regularization in the neuroimaging study (*α* = 0.1, *λ* = 0.0020, *MPSE* = 0.0023), and 29 features survived the regularization in the replication study (*α* = 0.6, *λ* = 0.0001, *MPSE* = 0.0062). In the motor task, a group of 25 features survived the regularization in the behavioral study (*α* = 0.2, *λ* = 0.0012, *MPSE* = 0.0039), 25 features survived the regularization in the neuroimaging study (*α* = 0.6, *λ* = 0.0001, *MPSE* = 0.0024) and 26 features survived the regularization in the replication study (*α* = 0.6, *λ* = 0.0001, *MPSE* = 0.0067). In the numerical task, 33 features survived the regularization in the behavioral study (*α* = 0.4, *λ* = 0.0001, *MPSE* = 0.0039), and 16 features survived the regularization in the replication study (*α* = 0.8, *λ* = 0.0013, *MPSE* = 0.0068). The list of surviving features appears in Supplementary Tables 4-5. See Supplementary Note 5 for the PCA dimensionality reduction strategy.

#### Relating choice dynamics to inconsistency levels (main task)

Following dimensionality reduction, we tested whether choice dynamics could predict inconsistency levels, using a subjects fixed-effects (random intercept) regression on motor components:

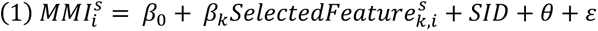

where *s* ∈ {1, … ,89}in the behavioral study, *s* ∈ {1, … ,42} in the neuroimaging study, and *s* ∈ {1, … ,89} in the replication study, is an index indicating subjects’ identity, *i* is an index indicating the trial number, *k* is an index indicating the mouse features that survived the elastic net regularization. *SID* is a subjects fixed-effect (random intercept) and *θ* is a vector of control variables including SV, budget set parameters (slope and endowment), RT and the CSI index (see above). We ran this model, with and without *θ*, on the data collected in each study using the mouse features from the risky-choice task. This analysis yielded that 18, 11 and 21 features (behavioral, neuroimaging and replication studies, respectively) correlated with MMI. See Supplementary Note 5 for the models used in the PCA strategy.

#### Leave-one-subject-out procedure

To obtain an out-of-sample prediction between mouse-tracking data and inconsistency levels, we repeated the dimensionality reduction procedure for *N* − 1 subjects, followed by estimation of regression coefficients for those *N* − 1 subjects (same model as in the previous paragraph). We then correlated the predicted trial-level inconsistency scores with the actual MMI scores. We repeated this process 89, 42, or 69 times (depending on the number of subjects in each sample – behavioral, neuroimaging, or replication studies), and reported the subject-specific correlation coefficients (Fig. 4e).

#### Relating motor output from the non-value motor tasks to choice inconsistency

To evaluate whether motion components unrelated to value modulation also correlated with inconsistency scores, we repeated the dimensionality reduction process on mouse features from the non-value motor tasks (in each of the two studies). We then used the independent motor components from the non-value tasks to predict MMI scores in the main risky-choice task in a subjects fixed-effects (random intercept) regression. Note that for the purpose of this analysis, in the behavioral and replication studies we could only use the subset of 51 trials that were common across the three tasks, whereas in the neuroimaging study all 75 trials were common across tasks and thus all 75 trials were used in this analysis (Fig. 4b).

In the elastic net strategy, we ran the following model:

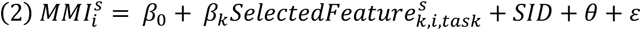

where *task* ∈ {*motor*, *numerical*} indicates whether the mouse feature was obtained in the numerical or motor tasks. All other notations are identical to equation (1). We ran this model, with and without *θ*, on the data collected in each study (separately) using mouse features from each of the non-value motor tasks (in separate models) as predictors of inconsistency levels. In the motor task 7, 6 and 9 features (behavioral, neuroimaging and replication studies, respectively) correlated with MMI, whereas in the numerical task 8 and 7 features correlated with MMI (behavioral and replication studies, respectively). We evaluated goodness of fit by examining the adj-R^2^ of each model (see Supplementary Note 5 for the models used in the PCA strategy).

#### fMRI data preprocessing

BrainVoyager QX (Brain Innovation) was used for image analysis, with additional analyses performed in MATLAB (MathWorks). Functional images were sinc-interpolated in time to adjust for staggered slice acquisition, corrected for any head movement by realigning all volumes to the first volume of the scanning session using six-parameter rigid body transformations, and de-trended and high-pass filtered to remove low-frequency drift in the fMRI signal. Spatial smoothing with a 6-mm FWHM Gaussian kernel was applied to the fMRI images. Images were then co-registered with each subject’s high-resolution anatomical scan and normalized using the Montreal Neurological Institute (MNI) template. All spatial transformations of the functional data used trilinear interpolation.

#### Functional ROIs

We focused our analysis on predefined hypothesis-driven regions of interest (ROIs). To maintain continuity with our previous work, all value-related brain regions (vmPFC, dACC, vStr and PCC) were defined in the same way as in Kurtz-David et al.^27^ In practice, the vStr and the vmPFC were defined using the masking in the canonical meta-analysis for value encoding by Bartra et al.^9^ For the dACC we drew a 12-mm sphere around the peak voxel reported by Kolling et al.^90^ who looked for positive neural correlates of foraging value, while controlling for RT and other task parameters in a RFX GLM. For the PCC, we simply used neurosynth.org meta-analyses masks with “PCC” as the search words.

We used V1 as a control region to rule out the possibility that SV and MMI correlated with a vast number of brain regions. We made use of the connectivity-based parcellation atlas by Fan et al.^91^, and drew 12-mm spheres around the peak voxels (one for each hemisphere) from their sub-region probabilistic voxel-mapping.

To define the motor brain regions, left-M1 and SMA, we used a degenerated version of the motor task that appeared in our previous study^27^ as an independent functional localizer. We estimated a general linear model (GLM) with one predictor that modeled RT using a boxcar epoch function convolved with the canonical hemodynamic response function (HRF), whose duration was equal to the RT of the trial. Six motion-correction parameters and the constant were included as nuisance regressors of no interest to account for head motion-related artifacts. We used a stringent Bonferroni correction as a statistical threshold, and defined any significant voxel with a positive correlation as either belonging to M1 or the SMA regions. Finally, to avoid overlapping between ROIs, we excluded from the SMA ROI any voxels that were common with the neighboring dACC ROI. See Supplementary Fig. 6 to see the resulting M1 and SMA functional ROIs.

#### Neural correlates of choice inconsistency and replication of the main finding in Kurtz-David et al

To identify the neural correlates of choice inconsistency in the main task, we ran exactly the same GLM regression that appeared in our previous study (see Kurtz-David et al. for further details)^27^. Model regression:

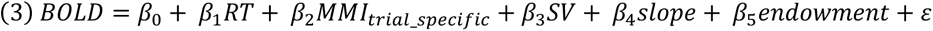

We tested the model on all our predefined value-related ROIs. Furthermore, we used the same model to examine neural correlates of choice inconsistency and value modulation in M1 and SMA.

Since a large portion of our sample (8 out of 42 subjects, 19%) was consistent with GARP, we wanted to increase statistical power for a whole-brain analysis. Hence, we also ran a more refined model, which only included trial-specific-MMI and a boxcar epoch function to model RT, both were entered for the trial duration and convolved with the canonical HRF. Six motion-correction parameters and the constant were included as regressors of no interest to account for motion-related artifacts.

#### Neural correlates of mouse features

To evaluate the neural correlates of motor output during task execution, we ran an RFX GLM with mouse features that survived the elastic net regularization and significantly correlated with MMI in both tasks of the neuroimaging study.

Namely, we used meanVel, MD and numFixations as our predictors of interest (Fig. 4c and Supplementary Tables 4-5). Additional predictors included budget sets parameters (slopes and endowments) and a boxcar epoch function to capture RT. All predictors were normalized, convolved with the HRF and entered for the trial duration. Six nuisance head movements regressors and a constant were included as well. We ran the model on each of the value and motor-related ROIs, and looked for differences in the neural representation of motor output between the value-based task compared with the motor task.

#### Statistical thresholds

All the GLMs in the paper were run at the voxel-level, and the results reported in Figs.6-7a present significant voxels that survived the following criteria: (a) ROI analyses - to increase the statistical power, the vast majority of the neuroimaging analyses focused on hypothesis-driven ROIs. In those analyses, in each ROI, we determined a statistical threshold of p<0.05 with at least six adjacent voxels that meeting this criterion. (b) Whole-brain analysis – the analysis presented in Fig, 6d is the only whole-brain analysis in the paper, aimed to strengthen the findings of Fig.6a. Here, we determined a more stringent threshold of p<0.005 with at least six adjacent voxels meeting this criterion. We further applied a permutation test with 1,000 iterations, and reported the significant voxels.

#### PPI

We performed a PPI analysis^92^ to examine the changes of task-related connectivity between value and motor brain regions as a function of inconsistency levels. The value related ROIs were selected as seeds (vmPFC, vStr, dACC and PCC) with two separate PPI analyses were done for each seed, one for each motor ROI (M1 and SMA, for a total of 8 models). The PPI regressors were thus generated by multiplying the demeaned BOLD time series of M1 or SMA with the trial-specific MMI (convolved with the canonical HRF). The other regressors that were included in the model were: motor regions’ demeaned BOLD time series, MMI, budget sets parameters (slopes and endowments) and a boxcar epoch function to capture RT. All predictors except the BOLD signal were convolved with the HRF and entered for the trial duration. We used the Bonferroni correction for multiple comparisons (8 models), and report significant results of the PPI regressors for a threshold of 0.00625.

#### Ruling out value modulation in the motor task

To show that our task manipulation worked, and that the additional non-value motor task indeed did not involve value modulation and solely examined motor output, we conducted two separate analyses.

First, we conducted a pseudo-SV analysis and treated the motor task in the neuroimaging study *as if* it actually was a value-based choice task. Given this thought experiment, we treated the coordinates of each mouse button-press at the end of the trial like a lottery chosen in the main risky-choice task. We then elicited subject-specific utility parameters and calculated the “trial-specific-SV” and “trial-specific-MMI” to be used as parametric regressors in a GLM with a design matrix identical to the GLM used to look for the neural correlates of choice inconsistency in the main risky-choice task (see above). We ran this model against the BOLD signal in all predefined value-related ROIs. Our main hypothesis in this analysis was to find null positive correlates between the pseudo-SV predictor and the BOLD signal in value-related brain regions, suggesting that the motor task did not result in value modulation processes in the brain (Fig. 6b).

Our second approach was to test whether the motor task induced a different type of decision-making mechanism, one that did not involve utility maximization, nor could it yield choice inconsistencies. We hypothesized that during task execution, subjects had to constantly determine whether they had reached the target coordinate. We therefore used the equal-time sampled trajectories (InterTime, see Supplementary Note 2) upon which we calculated the exact mouse-location in each TR using a linear interpolation. We then calculated for each such location the Euclidean distance between current location and target. The Euclidean distance between current location and target at each TR was used as a parametric regressor in an RFX GLM, normalized and convolved with the HRF. Additional predictors included the slope and Y-axis intersection of the linear graph, as well as an epoch function to capture RT (all normalized and convolved with the HRF). This model was tested against all the predefined value/decision-making ROIs to look for brain regions that tracked distance from target as a reinforcement-like signal.

### Data & code availability

The code used for the computation of MMI and the other inconsistency indices is available as an open-source code at https://github.com/persitzd/RP-Toolkit. The code package used to generate all mouse features, sample raw data and the code used to generate all the figures in the paper is available as an open source at https://github.com/djlevylab/trembling-hand-unraveled.

## Acknowledgements

We thank P. Glimcher and D. Persitz for thoughtful comments, D. Hasson for his help in running the behavioral study and O. Nofekh (R.I.P) for help with the inconsistency indices code package. This work was funded with a grant from the Israel Science Foundation (#954/17 for D.J.L) and the financial support of the Henry Crown Institute of Business Research in Israel.

## Author contributions

V.K.D and D.J.L conceived the study. D.J.L, A.H., and V.K.D designed the study. V.K.D collected the data of the behavioral and neuroimaging studies, and A.M. collected the data of the replication study. A.H. extracted the mouse features. V.K.D performed all other behavioral data analyses. V.K.D and A.M. performed the neuroimaging analyses. N.P helped with fMRI data preprocessing. V.K.D., A.M, A.H. & D.J.L wrote the manuscript.

## Notes

https://github.com/djlevylab/trembling-hand-unraveled

### Competing Interest Statement

The authors have declared no competing interest.

### Summary of Updates

Given the explorative manner of our analysis approach, we conducted a replication to strengthen our findings. All main findings in the paper were replicated. We further updated some of the analyses and edited the text throughout the paper.

