## Supplementary information for "The trembling hand unraveled: motor and valuation elements in the neural sources of choice inconsistency"

### Supplementary Notes

#### Supplementary Note 1. An English version of the instructions sheet of the non-value motor cognition tasks (behavioral study)

*We detail here the instructions from the additional motor cognition tasks from the behavioral study. Subjects received these instructions once they completed the main choice task. In the neuroimaging study, subjects received the instructions for the motor task at the beginning of the experiment together with the instructions for the main choice task. In addition, the maximal payoff for the motor task in the neuroimaging study was 20 NIS rather than 5 NIS, and show-up fee was 100 NIS rather than 25 NIS. Aside from specifics regarding the scanning procedure, instructions for the main choice task are identical to the instructions in Kurtz-David et al. (2019)[1].*

We will now ask you to complete two additional tasks. The order of the tasks will be chosen at random. Some subjects will first conduct TASK II and then move on to TASK III, and some subjects will first conduct TASK III, and then move on to TASK II.

**TASK II**

In this task, you will be presented with different graphs, similar to the linear graphs you saw in TASK I with one major change: there will be no numbers on the graphs, nor on the axes-grid. A black circle will be marked on each graph. In this task, you will have to move the mouse cursor from the axes-origin towards the clack circle, and left-click the mouse, once you reached the circle. You will have 6 seconds to reach the black circle. Try to be as precise as possible. After you left-click the mouse, your answer is submitted, and you will not be able to change it. If by the end of the 6 seconds, you have not submitted an answer, the computer will notify you with the following message: "Please note! In this round you did not left-click the mouse." Soon after you submit your answer, and/or at the end of the 6 seconds, you will move to a black screen. We shall ask you not to move the mouse during the black screen. Next, a new graph will appear on the screen, and you will be asked to reach a new circle. You will see a total of 51 such graphs.

**TASK III**

In this task, you will see various graphs with numbers on the axes-grid, similarly to the main task (TASK I) in the experiment. Here, you will see X & Y values (a coordinate) at the top of the screen. This coordinate is your target on the graph. You will have to move the mouse and reach the numeric coordinate as precisely and as soon as possible. After you reach the coordinate, left-click the mouse to submit your answer. For example, if at the top of the screen, you will see the values (20,50), you will have to scroll the mouse towards the coordinate (20,50). Note that the left number represents the X value, and that the right number represents the Y value. Similarly to TASK I, you will see at the top of the screen the current cursor position at any given moment. You will have 12 seconds to reach the numerical coordinate. Once you left-click the mouse, your answer is submitted, and you will not be able to change it. If by the end of the 12 seconds, you will not submit an answer, the computer will notify you with the following message: "Please note! In this round you did not left-click the mouse." Soon after you submit your answer, and/or at the end of the 12 seconds, you will move to a black screen. We shall ask you not to move the mouse during the black screen. Next, a new graph will appear on the screen, and you will be asked to reach a new coordinate target. You will see a total of 51 graphs. All the numerical targets in this task are points on the graph.

**Payoff for TASK II and TASK III**

You will receive an additional 5 NIS for participating in each of TASK II and TASK III (a total of 10 NIS). However, you might lose some of this amount, according to your performance in these tasks. The payoff structure in each task is the same, and goes as follows:

At the end of each task, one trial will be chosen at random. Each trial has an equal chance to be chosen. The computer will notify you what was your target in this trial, and will also show you your submitted coordinate (the point where you left-clicked the mouse). In both TASK II and TASK III, your target and choice will be represented by (X,Y) coordinates, whether you could see them during the task (as in TASK III) or not (as in TASK II). Soon after the computer will show you your payoff that will be determined according to your precision in the trial that was randomly chosen. The greater the distance between the target and choice, the larger is the amount that you will lose off your payoff. Therefore, we recommend you to be as precise as possible. The computer will also present you with the mathematical equation that calculates your distance from the target, and its impact on your payoff.

As mentioned, your maximal payoff for each of TASK II and III is 5 NIS. You can win this amount if you left-click *exactly* at your target coordinate.

Similarly to TASK I, if the computer randomly chooses a trial in which you did not submit a choice, then the computer will cast a random coordinate on the graph, and your payoff will be calculated according to this random coordinate.

To sum up, your total payoff will include four elements:

(a) 25 NIS participation fee
(b) The lottery results in TASK I, i.e. the monetary value of the product that was casted, X or Y, in the decision problem that was chosen at random.

(c) Your payoff, according to performance in TASK II, up to a maximum of 5 NIS
(d) Your payoff, according to performance in TASK III, up to a maximum of 5 NIS

**General instructions**

Please do not talk to anyone during the experiment. We ask all participants to remain quiet, up until the last participant completes the experiment.

**Thank you and good luck!**

#### Supplementary Note 2: Extended Features Description

The following passages detail all transformations of and calculations on each mouse trajectory of each trial in the experiment. Note that a mouse trajectory included the x and y coordinates of the mouse at every recorded time point throughout a trial. The units of the coordinates corresponded to the units of the graph, and so they were the values of the budgets at the mouse curser's current location, in tokens, which were converted to NIS at the end of the experiment.

##### Mouse Trajectory Modifications

First, the original mouse trajectory was saved as "**OrgTraj**". A new mouse coordinate was saved only whenever the mouse had moved, meaning the sampling rate of coordinates varied, providing an unevenly spaced time vector that coincided with the mouse coordinates of a trial. Since trials varied in the amount of motion during a trial, and trials differed in their length (also referred to as their reaction time), each trajectory had a different number of coordinates recorded. To even out the number of coordinates across trials, and to equalize the time intervals between each coordinate sample, we performed linear interpolation on both the time vector and on each of the x and y coordinates separately, which we saved as "**InterpTime**" and "**InterpOrg**", respectively. This was done using MATLAB's "interp1" function, and it resulted in 100 samples of coordinates per trial, and 100 timepoints per trial to match them. Due to the interpolation, these timepoints had a constant sampling rate, saved as "**SR**", but each trial had its own unique sampling rate since trials differed in their overall length. For example, a longer trial would mean a lower sampling rate to reach the final 100 samples, while a faster trial would mean a higher sampling rate to reach the evenly spaced 100 samples. Additionally, we wanted to create a version of the trajectory that had finer resolution and more samples per trials, so we conducted the same procedure but arriving at 1,000 final coordinates per trial (**"InterpOrgHighRes"**) and 1,000 timepoint (**"interpTimeHighRes"**), and a larger sampling rate ("**SRHighRes**"). Finally, we created a version of the trajectory, which was not sampled at a constant rate of time, but rather sampled equally in space. MATLAB's "interparc"[2] function enables to interpolate a set of points at fixed distances along some curve in space, by employing spline interpolation formulated in terms of differential equations that describe the path along the curve. This resulted in new mouse coordinates, saved as "**EqualTraj**", which were uniformly spaced along the same curve, rendering their time vector meaningless, but enabling more accurate calculations on the mouse trajectory's curve. However, since the "EqualTraj" version of the trajectory was intended for geometric calculation on the trajectory shape between the axes-origin to the budget line, we did not want this trajectory to include any additional mouse movements along the budget line itself. Therefore, this trajectory included only mouse coordinates from the beginning of a trial, in the origin, to the first arrival to the budget line, defined as reaching within 2 tokens of the budget line (“**Traj2Line**”). All other mouse coordinates after first reaching the budget line were discarded.

**Velocity Features**

These features relate to various aspects of the speed at which decisions were executed, which often fluctuated throughout the trial in manners that could elucidate the decision process. We started by measuring the Euclidean distance between each two consecutive points in “InterpOrg”, yielding the “**Seg_D_Progress**” vector. Next, the maximum progress made between two consecutive samples, which was the maximum of the previous vector, was saved as “**MaxProgress**”, and the time passed in the trial to reach this maximal progress was saved as “**Time2MaxProgress**”. The median of the vector was also saved as “**MedianProgress**“. The velocity at each timepoint was calculated by dividing distances in “**Seg_D_Progress**” by the differences of the “interpTime” vector. The feature “**VelOver10**” was the total duration spent in velocity which was over 10 tokens per second, calculated by multiplying the number of samples with velocity higher than 10, by the duration of a sampling interval. Then, the mean, median, and max velocities were also saved to “**meanVel**”, “**MedianVel**”, and “**MaxVel**”, respectively, while also extracting the time passed from the beginning of the trial up to reaching the max velocity, termed “**MaxVelTime**”. The feature “**EndofTrialVel**” was the mean velocity in the last 10 timepoints of velocity vector. Finally, a vector for “**Acceleration**” was created by dividing the differences in velocity again by the differences in “interpTime”. The mean acceleration was saved as “**meanAcc**” and the max acceleration as “**MaxAcc’**. Using MATLAB’s “findpeaks” function, with a minimal peak distance of 3, min height of 50 and a threshold of 1, we counted the number of peaks in the acceleration vector and saved the count as “N_Accs”.

**Complexity Features**

Complexity features attempt to assess how convoluted, tangled, and chaotic was the mouse trajectory, in contrast to a clean and smooth movement of the mouse throughout a trial. The “**angles”** of a trajectory were calculated as the absolute inverse tangent of the quotient between the x and y coordinate of each sample. The “**AngleSTD**” was then the standard deviation across all angles of a trial’s trajectory. Moreover, for a given trial, we calculated the sample entropy with a window of size 4 on each of the x and y times-series separately, using MATLAB's "sampen" function[3,4] ("**'XSampEn4'**"," **YSampEn4**"). Lastly, each time the subject changed the direction of the mouse on the x-axis, the derivative of the x axis time-series changed from positive to negative, or vice versa. So, using these changes in the derivative we counted the number of times the subject flipped directions in the x-axis ("**Xflips**") and the longest time between two consecutive flips ("**MaxXflip**"). To avoid counting small perturbation of the time-series that did not constitute a meaningful flip, we smoothed the time-series beforehand, using MATLAB "smooth" function with its default parameters. The same procedure was applied to the y-axis time-series, to obtain "**Yflips**" and "**MaxYflip**".

**Directedness Features**

Features that measure directedness try to capture how directly the subject moved to her desired bundle in a trial, versus a more curved or bended movement. Whether she moved the mouse straight from the origin to her chosen bundle, or whether she took detours, swerved, deviated, or deliberated as she moved. The "**EqualTraj**" version of the trajectory was entered into MATLAB's function "curvature"[5]. This function calculates the circumcenter for every triplet of neighboring coordinates along the trajectory. The "**curvatures"** vector is the unit vector in the direction from the center coordinate to the center of the circle, divided by the radius, at each triplet. Since we were not interested in the direction of the curves, but only in the degree of curveness at each triplet, we summed the norms of the curvature vector, to reach a final measure of "**curveness**" for a trial's trajectory. The "curvature" function also outputs the arc of the circle for each triplet, so we saved the maximum arc as another feature termed "**ArcLenMax**". Additionally, we calculated two well-established measures for mouse trajectories (same freeman refs), Maximum Deviation ("**MD"**) and Area Under Curve ("**AUC**"). The maximum deviation was calculated by drawing a line between the axes-origin to the chosen bundle, then finding the furthest point in the trajectory from this line (in terms of perpendicular distance to the line), and then taking its distance from the line as our measure. The area under the curve was trickier, as the layout of our experimental design was different to most common paradigms that use this measure. The classic calculation assumes the target choice is in fixed location (either left or right of the screen), and hence uses the "trapz" function to calculate the area between the trajectory and the x-axis, and then subtracts the triangle area between the straight line to the target and the x-axis to obtain the AUC. In our experiment, the target choice could be any coordinate on the budget line, and the trajectories often passed through the straight line to the target several times, rendering this technique imprecise. So, we had to transform the trajectory into a mask image that was 100X100 pixels, representing a grid of the task's graph. Each pixel that the mouse trajectory had passed through and the straight line between the origin and the chosen bundle were given the value one, while all other pixels were zero. This resulted in a closed object within the mask image, which was bounded by the straight line and the mouse trajectory. Then, using MATLAB's "imfill" and "imclose", we filled the object, turning all pixels within it to a value of 1. Finally, we summed the number of pixels with a value of 1 in the mask image, and divide it by the total amount of pixels, to obtain our AUC measure.

**Fixation Features**

Occasionally, a subject would pause their movement and stay adjacent to the same coordinate for a lengthy period. We suspected that these pauses in movement, which we termed "fixations", would reflect deliberation or uncertainty in the decision process. Thus, we extracted various features that capture different aspects of the mouse fixation points. The first fixation point can be considered as the point of origin, as subjects often delayed a while in the origin before initiating their movement. We calculated the number of samples that were lower than 2 tokens in both their x and y coordinates in the "InterpOrgHighRes" trajectory, in the first 30% of their trial, and multiplied that by the time interval in "SRHighRes", to receive the total time the subject spent in the origin. For any other fixation points, we divided the task's graph (from 0 to 100) to a 20X20 grid, such that each bin had a width of 5 tokens on each axis. Next, using MATLAB's "hist3" function, we counted how many samples of a trajectory belonged in each bin, and multiplied that by the time interval for each sample to obtain to total time spent in that bin. Any bin wherein the subject spent more than 0.2 seconds with their mouse, was considered a fixation point. So, we counted the number of fixation points ("**NumFixations**"), the average time spent in all fixation points ("**AveFixTime**"), the longest time spent in a fixation point ("**MaxFixTime**"), the Euclidean distance from the longest fixation point to the final choice ("**MaxFixChoiceDist**") and to the predicted choice by the subject-specific elicited utility parameters ("**MaxFixPredictDist**"), and finally, the number of fixations that were within a distance of 2 tokens around the budget line ("**numBudgetFixations"**). Beyond fixation points, we also calculated the time spent up to 10 tokens from the predicted bundle ("**TimeNearPredicted**"), in Euclidean distance, and the time spent within 2 tokens above or below the budget line ("**TimeBudgetLine**"), measured as the perpendicular distance to the line. Lastly, we measured how many samples were below the x-axis, where $y<0$, and multiplied by the time interval to obtain "**XTimeOutofBounds**" and did the same for samples beyond the y axis (x<0) to obtain “**YTimeOutofBounds**”, and for samples that exceeded northeast bound above the budget line (“**AboveLine**”).

#### Supplementary Note 3: Eliciting utility parameters and measuring trial-by-trial inconsistency levels in the main task

***The decision problem.*** In each trial, the subject is faced a visualization of a budget line. Each discrete $(x, y)$ point on the budget line corresponds to a lottery with a 50% chance of winning the tokens allocated to the X-axis coordinate and a 50% chance of winning the tokens allocated to the Y-axis coordinate. Thus, the budget line describes all the attainable allocations to *X* and Y on a two-dimensional graph (bundles), such that its slope corresponds to the substitution – or *price ratio* – between X and Y. Formally – $p_1 x + p_2 y = E$, where $E$ is the total endowment to be spent on $X$ and $Y$, and $p_1$ and $p_2$ are the prices of allocating the endowment to either account. Thus, $\frac{p_2}{p_1}$ is the price ratio between the two accounts, and is graphically represented by the slope of the budget line.

***The General Axiom of Revealed Preference (GARP).*** Consider a finite dataset $\mathbf{D} = \{(\mathbf{p}^i, \mathbf{x}^i)\}_{i=1}^{n}$ , where $\mathbf{x}^i$ is the subject's chosen bundle at prices $\mathbf{p}^i$. Bundle $\mathbf{x}^i$ is

1. Directly revealed preferred to another bundle $\mathbf{x}$, denoted $\mathbf{x}^i R^0 \mathbf{x}$, if $\mathbf{p}^i \mathbf{x}^i \geq \mathbf{p}^i \mathbf{x}$.
2. Strictly directly revealed preferred to bundle $\mathbf{x}$, denoted $\mathbf{x}^i P^0 \mathbf{x}$, if $\mathbf{p}^i \mathbf{x}^i > \mathbf{p}^i \mathbf{x}$.
3. Revealed preferred to bundle $\mathbf{x}$, denoted $\mathbf{x}^i R \mathbf{x}$, if there exists a sequence of observed bundles $(\mathbf{x}^j, \mathbf{x}^k, \dots, \mathbf{x}^m)$, that are directly revealed preferred to one another, $\mathbf{x}^i R^0 \mathbf{x}^j, \mathbf{x}^j R^0 \mathbf{x}^k, \dots, \mathbf{x}^m R^0 \mathbf{x}$. Relation $R$ is therefore the transitive closure of the directly revealed preferred relation.

$\mathbf{D}$ satisfies the General Axiom of Revealed Preference (GARP), if every pair of observed bundles, $\mathbf{x}^i R \mathbf{x}^j$, implies $\neg(\mathbf{x}^j P^0 \mathbf{x}^i)$. We say a subject is consistent *iff* she satisfies GARP. We say that a utility function $u(\mathbf{x})$ rationalizes **D** if $\mathbf{x}^i R^0 \mathbf{x}$ implies $u(\mathbf{x}^i) \geq u(\mathbf{x})$. According to Afriat Theorem[6,7], there exists a well-behaved utility function (continuous, monotone and concave) that rationalizes the data *iff* the subject satisfies GARP. Otherwise, a strict cycle of choices exists and we say that **D** violates GARP. By Afriat's Theorem, if the data set $\mathbf{D}$ does not satisfy GARP, then the subject cannot be described as a non-satiated utility maximizer and is therefore said to be inconsistent[1,2].

***Aggregate inconsistency indices.*** We use three different indices to measure subjects' inconsistency levels. The first is simply counting the *number of GARP violations*, which quantifies the frequency of inconsistencies (Supplementary Fig. 7a). We use Afriat Index (AI)[8] and the Money Metric Index (MMI)[9] to evaluate the intensity of inconsistencies. *AI* measures the maximal adjustment to the budget sets needed to remove all violations. It can be interpreted as the subject's waste of income due to inconsistency (Supplementary

Fig. 7a). *MMI:* consider the continuous and non-satiated utility function $u(\cdot)$ as representing the preferences of the subject. $u(\cdot)$ induces a complete preference ranking for each bundle presented to the subject. At the same time, the subject provides a partial ranking of bundles by the principle of revealed preference. If these two rankings are compatible for every choice made by the subject, then $u(\cdot)$ rationalizes the dataset. If, however, these two rankings are incompatible, then according to $u(\cdot)$, some feasible bundles are ranked higher by $u(\cdot)$ than the bundle chosen by the subject. For every observation, the incompatibility between the two rankings can be measured by the minimal expenditure (a shift of the budget line towards the axes-origin) such that the adjusted budget set does not include any bundle that is strictly preferred over $\mathbf{x}^i$ according to $u(\cdot)$. Formally, given the budget set prices $\mathbf{p}^i$, the money metric $m(\mathbf{x}^i, \mathbf{p}^i, u)$ for observation $i$ is the minimal expenditure required for the dataset to include a bundle $\mathbf{y}$ such that – $u(\mathbf{y}) \geq u(\mathbf{x}^i)$: $m(\mathbf{x}^i, \mathbf{p}^i, u) = \min_{u(\mathbf{y}) \geq u(\mathbf{x}^i)} \mathbf{p}^i\mathbf{y}.$

We aggregate the adjustments for all observations using an average sum-of-squares (ANG(SSQ)) aggregator function. We examine all utility functions in the family of Disappointment Aversion utility functions (see below)[10] and look for the one with the smallest incompatibility with the dataset. Therefore, the computation of MMI yields two measures: (a) the computation of Aggregate MMI*;* (b) elicitation of subject-specific utility function parameters. See Supplementary Fig. 7b for a visualization of the index. For further details see Halevy et al. (2018)[9] and Supplementary Note 1 in Kurtz-David et al. (2019)[1].

***Trial-specific index.*** We implement a Leave-one-out procedure on the MMI index to yield a trial-level inconsistency index.[11] Let $\varepsilon_D$ be an aggregate inconsistency index of dataset $\mathbf{D}$. Let $\mathbf{D}_{-i}$ be a subset of $\mathbf{D}$ that includes all $n-1$ trials but the $i^{th}$ observation. Let $\varepsilon_{D_{-i}}$ be the aggregate inconsistency index of $\mathbf{D}_{-i}$, and let $(\varepsilon_D - \varepsilon_{D_{-i}})$ be the trial-specific inconsistency index of trial $i$.

***Parametric Utility.*** For parametric family of utility functions we use the Disappointment Aversion model with CRRA functional form[10], as it includes many well-known types of preferences in the context of risk[12,13] and has become the common practice for utility recovery for this task[1,9,12,14]. Formally,

$$(\text{S.}1)\ \text{SV}(x_1^{\text{i}}, x_2^{\text{i}}) = \gamma\omega(\max\{x_1^{\text{i}}, x_2^{\text{i}}\}) + (1-\gamma)\omega(\min\{x_1^{\text{i}}, x_2^{\text{i}}\}), \qquad \text{(DA)}$$

$$(S.2)\ \gamma = \frac{1}{2+\beta}, \qquad -1 \leq \beta < \infty$$

$$(\mathrm{S.3})\ \omega(\mathrm{z}) = \begin{cases} \dfrac{\mathrm{z}^{1-\rho}}{1-\rho}, & \rho \geq 0 \\ \ln(\mathrm{z}), & \rho = 1 \end{cases} \qquad \text{(CRRA)}$$

where $\gamma$ is the weight of the better outcome, and $\omega$ is a CRRA utility index with a relative risk aversion parameter $\rho$. When $\beta = 0$ this is the common Expected Utility function with parameter $\rho$i. If, in addition $\rho = 0$, it is Expected Value and when $\rho = 1$ it is the Cobb-Douglas with equal exponents. See Kurtz-David et al. (2019)[1] for details on the different behavioral types the DA function represents.

To use subjective value (SV) as a parametric regressor in our analysis, we calculated the value of the Disappointment Aversion model with CRRA functional form at the chosen bundle $(x, \mathrm{y})$ in each trial $i$, using the subject's recovered parameters, $\beta$ and $\rho$.

The code-package used to extract utility parameters and compute all inconsistency indices is available in https://github.com/persitzd/RP-Toolkit.

#### Supplementary Note 4: Descriptive results from mouse tracking

To reveal choice dynamics with and without value modulation, in both studies, we recorded mouse trajectories of motion pathways in all three tasks. Supplementary Fig. 2a displays example trajectories of one subject from all three tasks of the same linear line (similar trial) in the behavioral study. Supplementary Fig. 2b displays example trajectories of another subject from the MRI study. In both Supplementary Fig, 2a and 2b, the left panel presents an example where the trajectories across tasks have clear commonalities, implying that basic motor traits shaped choice execution both in the value-based task and in the motor cognition tasks. The right panel, on the other hand, shows an example where trajectories across tasks are quite different from one another. Supplementary Fig 2c-d present the corresponding velocity profiles for the trials in Supplementary Fig.2a-b, which constitute the second dimension in trajectories analysis in addition to the moment-by-moment mouse-loci on the axes grid. In the behavioral study, smooth trials have one major segment of acceleration and deceleration, which usually characterizes smooth reaching routes[15]. On the other hand, in the MRI study, we trace two such segments, probably due to the fragmented-movement that characterizes motion with the MRI-compatible trackball. Note, however, the synchrony in velocity profiles across the main and motor tasks, which shows, yet again, the similar characteristics in trajectories from the same trial across the different tasks. Not surprisingly, in trials with complex dissimilar pathways, like the ones presented in the right panels of Supplementary Fig. 2a-b, we identify several segments of acceleration and deceleration within a trial (right panels of Supplementary Fig. 2c-d). Supplementary Fig. 2e-f presents the mean velocity profiles across all trials and subjects. On average, subjects were faster in the behavioral study (Supplementary Fig. 2e), probably due to easier task execution when using a regular computer-mouse compared with the MRI-compatible trackball. Furthermore, subjects were faster in the non-value motor tasks than in the main value-based task. We relate this to the fact that the non-value tasks did not require subjects to make a value-based decision, and targets were predefined. Overall, the commonalities and similarities in mouse-loci and in velocity profiles indicate that motor output is comparable across tasks in both the behavioral and the neuroimaging studies.

#### Supplementary Note 5: PCA strategy

##### *Dimensionality reduction via PCA*

To check the robustness of the results, we also employed an unsupervised algorithm for reducing the dimensionality of the data. To this end, we conducted a Principal Component Analysis (PCA) on the mouse features. Importantly, this method preserves all the mouse features (as opposed to the elastic net), but at the expense of the interpretability of the components. We kept the first ten components (in each of the two studies), which together accounted for 76.6% (behavioral study), 77.3% (neuroimaging study) and 78.2% (replication study) of the variance in the data. Following dimensionality reduction, we tested whether choice dynamics could predict inconsistency levels, using a subjects' fixed-effects regression on mouse features' PCs:

$$(S.4)\; MMI_i^s = \delta_0 + \delta_j PC_j + SID + \theta + \varepsilon$$

where $s \in \{1, \dots, 89\}$ in the behavioral study, $s \in \{1, \dots, 42\}$ in the neuroimaging study, and $s \in \{1, \dots, 69\}$ in the replications tudy, is an index indicating subjects' identity, $i$ is an index indicating the trial number, $PC_j \in \{1, \dots, 10\}$ are the first ten components from the PCA analysis. $SID$ is a subjects-fixed effect (random intercept) and $\theta$ is a vector of control variables including SV, budget set parameters, RT and the CSI index. We ran this model, with and without $\theta$, on the data collected in each study using the mouse features from the risky-choice task

##### *PCs from the value-based task account for inconsistency levels*

We found very similar results to the elastic-net approach (Supplementary Fig. 5). In the behavioral study, we found that nine out of the first ten PCs correlated with inconsistency levels (PCs1-6 and PCs 8-10, Supplementary Fig. 5a, left panel, p<0.05 in a subjects fixed-effects regression). Similarly, in the neuroimaging study, we found that the four out of the first 10 principal components correlated with inconsistency levels (PCs 6-9, Supplementary Fig. 5a, right panel, p<0.05 in a subjects fixed-effects regression). Likewise, in the replication study we found that seven out of the first 10 principal components correlated with inconsistency levels (PC 1, PCs 3-4, PCs 6-8, and PC 10). Similarly to the elastic net approach, we repeated the leave-one-subject-out analysis, and found a significant positive correlation between predicted and actual inconsistency scores for all subjects in the behavioral study, for 39 of 42 subjects in the neuroimaging study, and for 67 of 69 subjects in the replication study (p<0.0001, Behavioral study: t(88)= 29.971, CI = [0.3620,inf], median correlation r=0.3797; Neuroimaging study: t(41)= 7.727, CI = [0.1351,inf], median correlation r=0.1470; Replication study: t(68)=16.1498,

CI=[0.3508;inf], median correlation r=0.3882). One-sided one-sample t-test, Supplementary Fig. 5c). Goodness-of-fit measures show that task-execution mouse PCs accounted for 5.0% (behavioral study, figure shows results for the common trials across tasks. The results remain robust when including all trials from the risky-choice task), 3.2% (neuroimaging study), and 4.4% (replication study) of the variance of inconsistency scores (Supplementary Fig. 5b). This indicates, yet again, that mouse-tracking data could predict inconsistency levels in and out-of-sample.

***PCs from the non-value motor tasks also capture choice inconsistency***

To evaluate whether motion components unrelated to value modulation also correlate with inconsistency scores, we repeated the dimensionality reduction process on mouse features from the non-value motor tasks (in each of the two studies). We then used PC1-10 from the motor tasks to predict MMI scores in the main risky-choice task in a subjects' fixed-effects regression. Concretely, we ran the following model:

$$(S.5)\; MMI_i^s = \delta_0 + \delta_{j,task} PC_{j,task} + SID + \theta + \varepsilon$$

Where $PC_j,\; j \in \{1,2,\dots,10\}$ are the first ten components from the PCA analysis and $task \in \{motor, numerical\}$ indicates whether the mouse feature was obtained in the numerical or motor tasks. All other elements in the model are identical to equation (2) in the Main Text. Finally, here too we evaluate goodness of fit by examining the adj-$R^2$ of each model.

We found that principal components from the non-value tasks captured variance in actual MMI scores, as they captured 1.7%-4.5% of the variability in inconsistency scores (Supplementary Fig. 5b). Similarly to the analysis reported in the Main Text (using the Elastic Net strategy), principal components from the non-value tasks provided a significant cross-tasks prediction for MMI scores when using regression coefficients from the main task (p<0.0001, behavioral study: r=0.4782, CI=[0.4548;0.501], and r=0.501, CI=[0.4785; 0.5238], motor and numerical tasks, respectively. Neuroimaging study: r=0.4418, CI=[0.4122,0.4706]. Replication study: r=0.6194, CI=[0.5972;0.6407], and r=0.6135, CI=[0.5913;0.6348], motor and numerical tasks, respectively).

#### Supplementary Note 6: Choice simplicity index (CSI)

To measure choice difficulty in our task, we introduced the Choice Simplicity Index (CSI) in our previous study[1]. CSI takes into account the continuity of the budget line (as opposed to binary choice tasks) and subject-specific preferences.

To compute CSI, we discretized each budget line into 1,000 possible bundles. We then calculated the subjective value $v_{i,b,s}$ of each bundle $i$ along each budget set $b$ for each subject $s$ using the parameters elicited by the MMI. Next, we subtracted $v_{i,b,s}$ from the maximal subjective value of that budget line to obtain a $\Delta V_{i,b,s}$ measure for each bundle along the budget set. This procedure is equivalent to subtracting the difference between two options in a binary choice design. We then averaged all the $\Delta V_{i,b,s}$ across the 1,000 bundles, and normalized the result by the endowment of the given budget set to be able to compare across trials. Formally, denote $V_{b,s} = max_{i \in [1,\ldots,1{,}000]} v_{i,b,s}$. Then:

$$(S.5)\ \text{Choice Simplicity}_{b,s} = \frac{\frac{\sum_{i=1}^{N}[V_{b,s} - v_{i,b,s}]}{N}}{\text{Endowment}_b}$$

where $i \in [1,\ldots,1{,}000]$ is the bundle (N=1000); $b \in [1,\ldots,150]$ ($b \in [1,\ldots,75]$ is the decision problems in the behavioral and replication studies (neuroimaging study), and $s \in [1,\ldots,89]$ ($s \in [1,\ldots,42]$ or $s \in [1,\ldots,69]$) is the subject in the behavioral study (neuroimaging or replication studies).

Note that the higher is our index, the easier it is to make a choice, suggesting that trials with index values closer to 0, are the more difficult choices. For further details, see Kurtz-David et al.[1]

### Supplementary Figures

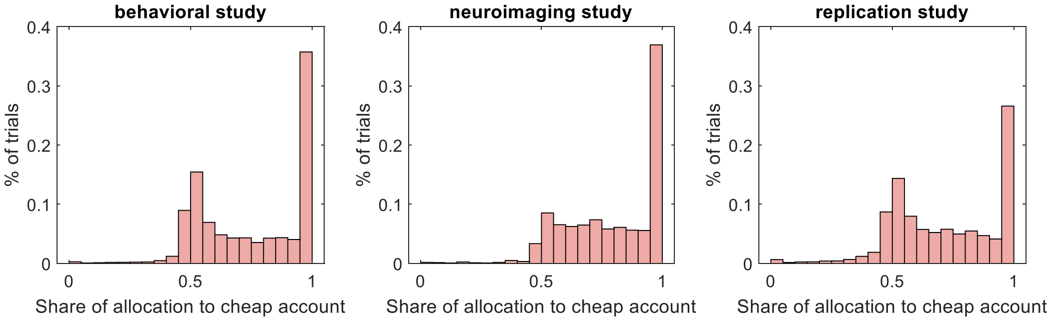


**Supplementary Figure 1 | distributions of trials by the share allocated to the cheap account (x or y).** *Left* – behavioral study; *middle* – neuroimaging study; *right* – replication study. When slope is> 1 (in absolute value), then it is more beneficial to allocate money to the Y-axis and it is considered the cheap account - and vis-à-vis - when the slope<=1 (in absolute value), then it is more beneficial to allocate more money to the X-axis, and it is considered the cheap account. As X & Y are symmetrical, allocating less than 50% of endowment to the cheap account corresponds to violating FOSD. The share of trials with FOSD violations was 11.79% in the behavioral study, 4.98% in the neuroimaging study, and 14.45% in the replication study. If we allow some sensitivity around the focal 50%-share point (the intersection between the budget set and a 45-degrees line from the axes origin) and examine only the share of trials where subjects allocated less than 45% to the cheap account, then we find that only 2.84% of trials in the behavioral study, 1.68% in the neuroimaging study, and 5.77% in the replication study, violated FOSD. This suggests that subjects understood the task.

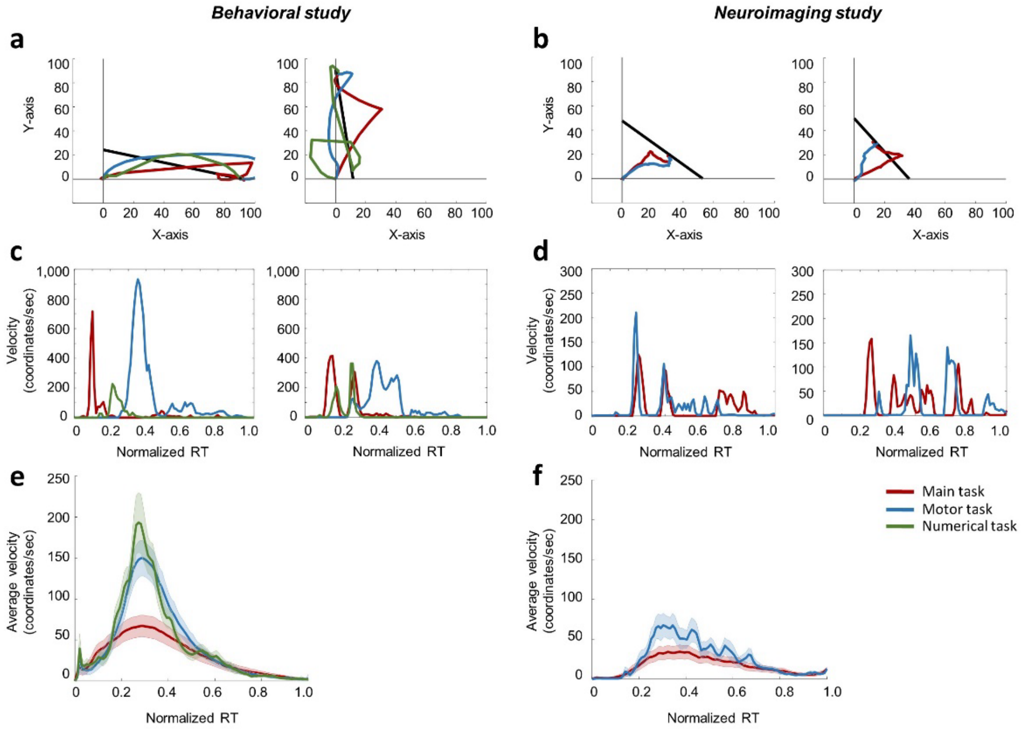


**Supplementary Figure 2 | Mouse tracking.** (a) Two trajectories from two different trials from the same subject (sub. 217) across the different tasks, behavioral study. *Left* – a trial with similar trajectories across tasks (trial 23). *Right* – a trial with dis-similar trajectories across tasks (trial 75). (b) Same as (a), but from a subject in the MRI study (trials 10 and 27 from subject 113). (c-d) Velocity profiles for the same trials presented in (a-b), respectively. We normalized RT to a relative scale. (e-f) Average velocity profiles for all subjects in the sample. We normalized RT to a relative scale. Shaded error bar represent standard errors. (e) Behavioral study. (f) MRI study.

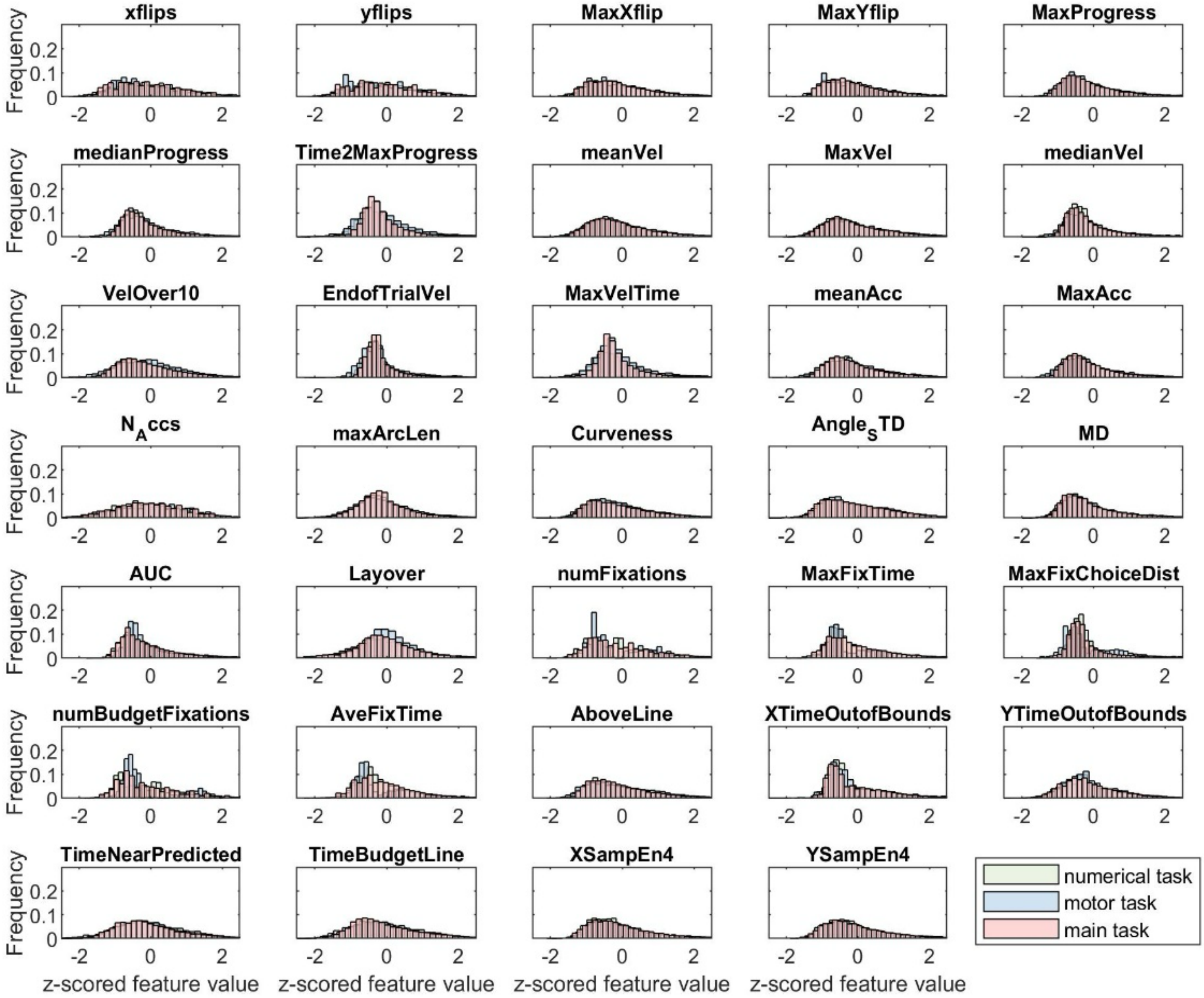


**Supplementary Figure 3 | Distribution of features values after preprocessing, behavioral study** (z-scoring, removing nan and extreme values).

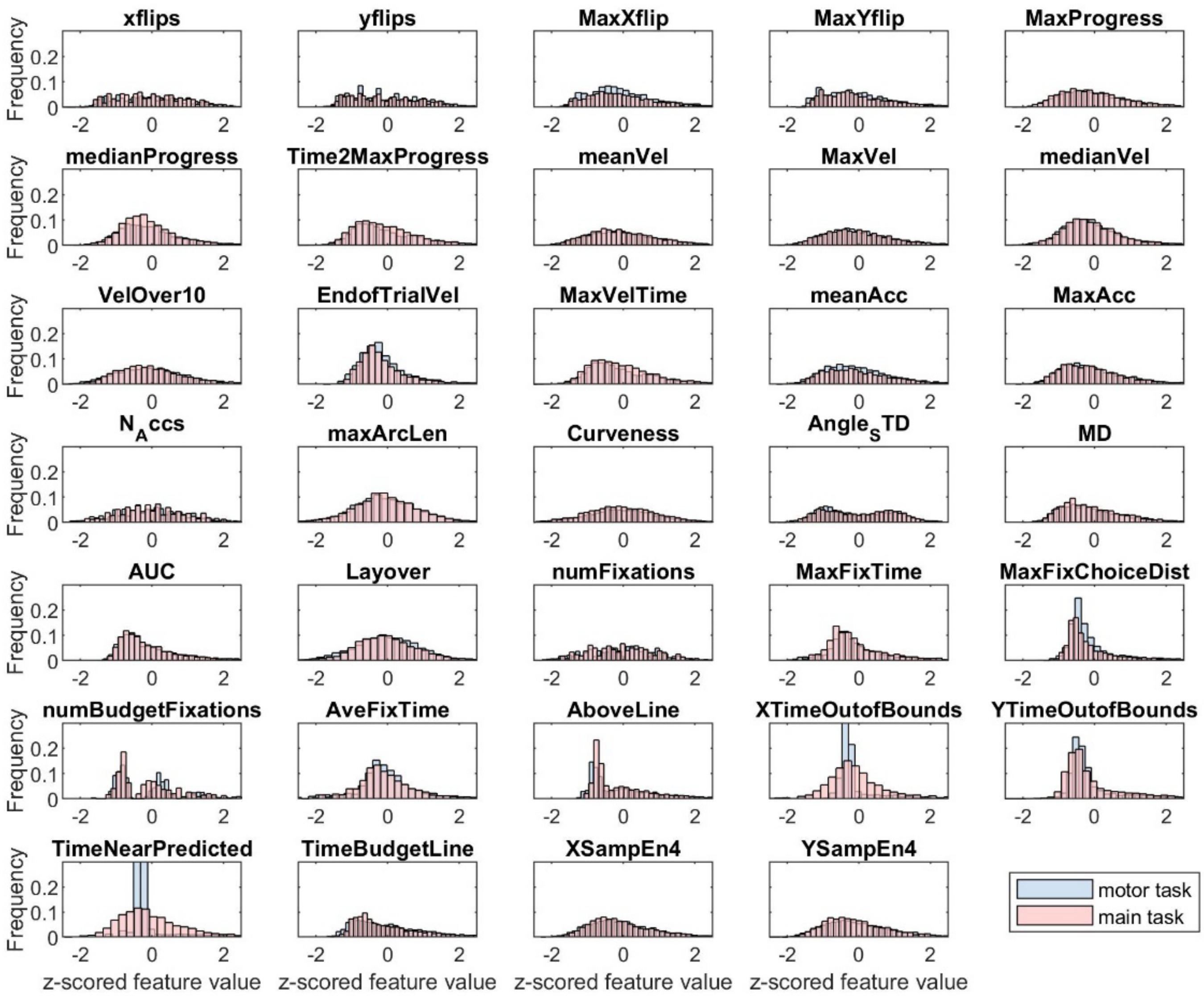


**Supplementary Figure 4 | Distribution of features values after preprocessing, MRI study** (z-scoring, removing nan and extreme values).

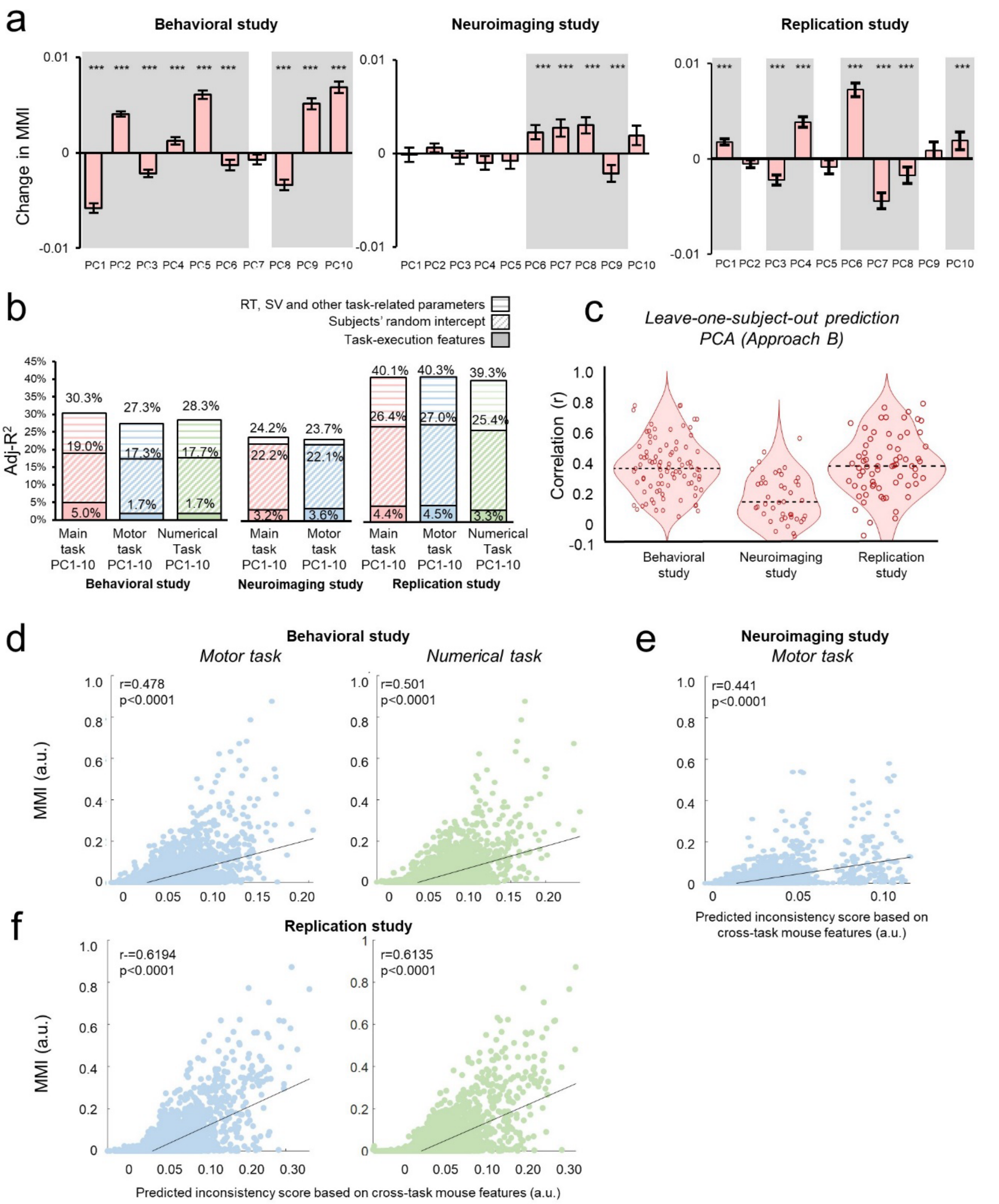


**Supplementary figure 5 | Mouse features and inconsistency levels, Approach B (PCA).** (a) Same analysis as in Fig. 4c, using Approach B – PCA. Mouse features in the main task correlated with inconsistency levels in both studies. Shaded area indicate a significant predictor (p<0.05) in a subjects fixed-effects regression (see Supplementary Note 6). *Left*: behavioral study. *Middle:* neuroimaging study. *Right*: replication study. (b) Adj-$R^2$ from regressions that included PCs from the main task, compared with regressions that included PCs from the non-value tasks. Solid fill indicates regressions, which only included task-execution mouse features. Diagonal striped fill indicates subjects' random intercept. Horizontal striped fill indicates additional explained variance from SV and other task-related parameters. *Left*: behavioral study, *middle:* neuroimaging study, *right:* replication study. For the behavioral and replication studies, the figure includes only the 51 trials that were common across all three tasks. (c) Leave-one-subject-out prediction. Distribution of correlation coefficients between predicted inconsistency levels based on PC1s from the main task and actual MMI scores. *Behavioral study:* median=0.3797, min=0.0980, max=0.7949, std=0.1582; *neuroimaging study:* median=0.142, min=-0.065, max=0.541, std=0.145. *Replication study:* median=0.382, min=-

0.109, max=0.7963, std=0.201. Dashed black line indicates median. Scatter shows individual correlation coefficients (behavioral study: N=89, neuroimaging study: N=42, replication study: N=69.). (d) Same analysis as in Fig. 5b-c, using Approach B – PCA. Obtained correlations between predicted and actual MMI. Behavioral study. Each dot in the scatter relates to a different trial. predictions based on mouse features from the behavioral study (N motor task = 4,279, N numerical task = 4,192). (e) Predictions based on mouse features from the neuroimaging study (N=2,915). (f) Predictions based on mouse features from the replication study (N motor task = 3,085, N numerical task = 3,160).

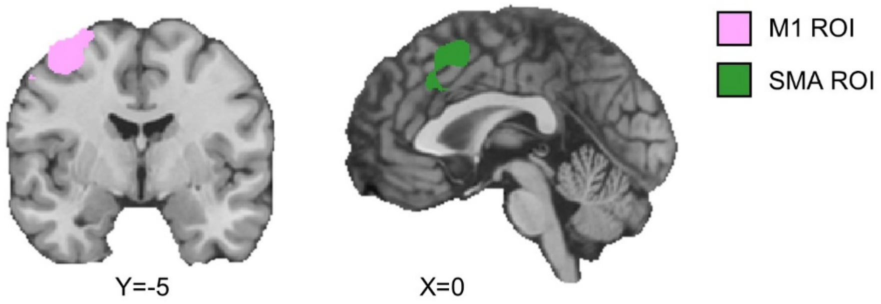


**Supplementary figure 6 | M1 and SMA functional ROIs.** We used a degenerated version of the motor task that appeared in Kurtz-David et al. (2019) as an independent functional localizer for m1 and SMA. Whole-brain RFX GLM, n = 22, q(Bonferroni)<0.05. Model regression: $BOLD = \beta 0 + \beta 1\, RT + \varepsilon$. We defined any significant voxel that correlated with the boxcar function as either belonging to m1 (left) or the SMA (right) regions. To avoid overlap between ROIs, we excluded from the SMA ROI any voxels that were common with the neighboring dACC ROI.

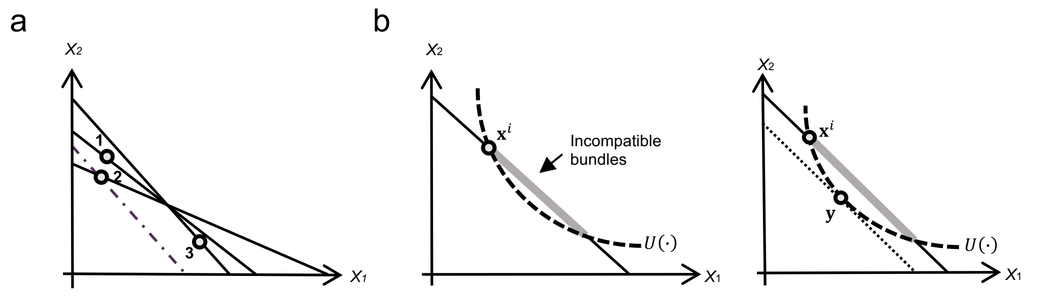


**Supplementary Fig. 7 |** An illustration of the inconsistency indices. (a) A dataset with choices on three budget lines. *Bundle 3* is reveled preferred to *Bundle 1* and *Bundle* 2, but at the same time *Bundle 1* and *Bundle 2* are reveled preferred to *Bundle* 3. These choices create a choice cycle with three GARP violations. The dashed line indicates Afriat Index (*AI*) and shows the biggest adjustment to the budget lines required to remove all the violations. (b) MMI. *Left:* The utility function $U(\cdot)$ ranks some bundles on the budget line higher than the ranking provided by the subject, who chose $\boldsymbol{x^i}$ as their most desired bundle. *Right:* We resolve this incompatibility by shifting the budget line towards bundle $\boldsymbol{y}$ (dotted line)**.** On the adjusted budget line, $y$ is the highest ranked bundle by $U(\cdot)$.

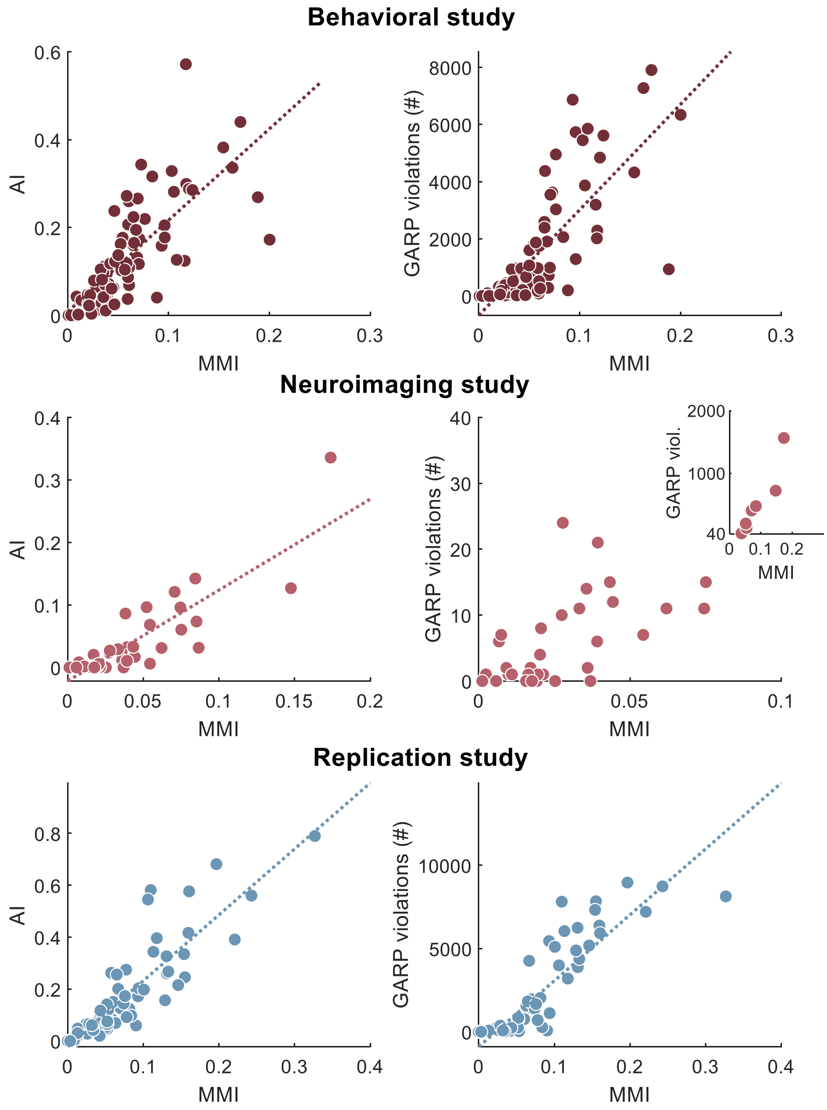


**Supplementary figure 8 | Correlations between inconsistency indices.** Scatterplots show correlations between inconsistency indices: upper panel shows the correlations between MMI and AI (r=0.7608, left) and the number of GARP violations (r=0.722, right) in the behavioral study, the middle panel shows the same results for the neuroimaging study (r=0.8618 and r=0.7989, respectively). The bottom panel shows the results for the replication study (r=0.8723 and r=0.8797, respectively). Dashed line indicates the least squares fit.

### Supplementary Tables

#### Supplementary Table 1 | Budget sets' parameters, behavioral study

| # | Slope ($p_y/p_x$) | Y intersection | # | Slope ($p_y/p_x$) | Y intersection | # | Slope ($p_y/p_x$) | Y intersection |
|---|---|---|---|---|---|---|---|---|
| 1 | -0.74 | 55.01 | 51 | -1.00 | 83.80 | 101 | -0.42 | 27.90 |
| 2 | -0.23 | 19.60 | 52 | -0.62 | 44.23 | 102 | -2.28 | 92.73 |
| 3 | -0.50 | 42.56 | 53 | -0.90 | 47.65 | 103 | -0.37 | 33.28 |
| 4 | -1.88 | 98.70 | 54 | -0.89 | 51.79 | 104 | -0.25 | 24.05 |
| 5 | -2.54 | 53.16 | 55 | -7.42 | 87.86 | 105 | -0.22 | 18.81 |
| 6 | -1.16 | 97.26 | 56 | -3.59 | 89.01 | 106 | -3.79 | 98.16 |
| 7 | -0.56 | 44.56 | 57 | -2.35 | 51.47 | 107 | -1.54 | 60.68 |
| 8 | -7.96 | 65.39 | 58 | -0.15 | 10.80 | 108 | -0.25 | 16.58 |
| 9 | -1.67 | 95.10 | 59 | -0.21 | 11.18 | 109 | -0.18 | 17.76 |
| 10 | -0.33 | 27.67 | 60 | -2.84 | 63.98 | 110 | -0.45 | 39.88 |
| 11 | -1.38 | 50.11 | 61 | -0.22 | 18.83 | 111 | -0.85 | 48.91 |
| 12 | -1.51 | 56.35 | 62 | -0.58 | 30.41 | 112 | -0.68 | 61.31 |
| 13 | -0.19 | 18.36 | 63 | -1.11 | 75.59 | 113 | -0.24 | 22.73 |
| 14 | -0.17 | 11.35 | 64 | -1.61 | 55.80 | 114 | -0.47 | 28.58 |
| 15 | -1.16 | 99.63 | 65 | -0.41 | 24.15 | 115 | -0.62 | 43.28 |
| 16 | -0.32 | 24.41 | 66 | -0.10 | 9.13 | 116 | -0.87 | 53.76 |
| 17 | -1.24 | 54.26 | 67 | -0.14 | 8.67 | 117 | -0.47 | 44.39 |
| 18 | -0.61 | 44.78 | 68 | -0.25 | 19.68 | 118 | -0.39 | 37.20 |
| 19 | -1.15 | 86.24 | 69 | -1.60 | 83.55 | 119 | -1.40 | 62.60 |
| 20 | -0.21 | 20.88 | 70 | -1.82 | 96.68 | 120 | -0.12 | 7.50 |
| 21 | -0.19 | 15.86 | 71 | -0.81 | 48.28 | 121 | -0.37 | 18.67 |
| 22 | -2.77 | 54.84 | 72 | -7.75 | 57.36 | 122 | -0.68 | 48.88 |
| 23 | -0.59 | 49.72 | 73 | -3.70 | 82.84 | 123 | -1.40 | 53.01 |
| 24 | -1.10 | 89.02 | 74 | -0.97 | 63.01 | 124 | -7.80 | 90.82 |
| 25 | -3.56 | 85.72 | 75 | -0.63 | 37.93 | 125 | -0.61 | 41.46 |
| 26 | -1.15 | 70.37 | 76 | -1.19 | 89.50 | 126 | -0.32 | 18.91 |
| 27 | -0.66 | 46.53 | 77 | -1.87 | 82.54 | 127 | -0.53 | 47.56 |
| 28 | -0.11 | 6.07 | 78 | -1.05 | 93.30 | 128 | -0.12 | 6.83 |
| 29 | -0.42 | 21.91 | 79 | -0.28 | 19.82 | 129 | -0.54 | 41.60 |
| 30 | -0.25 | 19.98 | 80 | -0.40 | 26.95 | 130 | -0.25 | 16.35 |
| 31 | -0.46 | 32.99 | 81 | -8.39 | 66.90 | 131 | -1.06 | 74.27 |
| 32 | -1.18 | 93.16 | 82 | -0.26 | 24.54 | 132 | -0.40 | 32.47 |
| 33 | -1.56 | 81.18 | 83 | -0.99 | 85.46 | 133 | -0.27 | 26.06 |
| 34 | -2.31 | 73.22 | 84 | -0.15 | 11.78 | 134 | -0.63 | 50.85 |
| 35 | -0.73 | 56.40 | 85 | -0.39 | 27.97 | 135 | -4.46 | 91.56 |
| 36 | -1.78 | 92.64 | 86 | -8.04 | 93.70 | 136 | -0.26 | 14.18 |
| 37 | -1.54 | 85.23 | 87 | -0.19 | 17.73 | 137 | -1.79 | 91.90 |
| 38 | -2.91 | 82.66 | 88 | -0.32 | 19.20 | 138 | -0.84 | 62.64 |
| 39 | -2.18 | 63.18 | 89 | -0.39 | 21.62 | 139 | -1.43 | 97.58 |
| 40 | -3.65 | 70.37 | 90 | -0.13 | 9.27 | 140 | -0.40 | 23.75 |
| 41 | -0.70 | 46.01 | 91 | -2.25 | 92.06 | 141 | -0.52 | 46.43 |
| 42 | -1.37 | 69.72 | 92 | -0.25 | 23.92 | 142 | -0.40 | 30.30 |
| 43 | -0.35 | 31.17 | 93 | -3.80 | 61.03 | 143 | -0.43 | 34.23 |
| 44 | -0.75 | 60.24 | 94 | -0.95 | 85.38 | 144 | -0.57 | 51.98 |
| 45 | -0.47 | 25.88 | 95 | -4.39 | 54.83 | 145 | -0.32 | 30.45 |
| 46 | -2.59 | 64.18 | 96 | -3.18 | 85.77 | 146 | -0.90 | 49.37 |
| 47 | -1.32 | 58.36 | 97 | -0.20 | 19.43 | 147 | -0.26 | 24.33 |
| 48 | -0.54 | 45.83 | 98 | -0.36 | 32.29 | 148 | -0.41 | 39.22 |
| 49 | -1.64 | 80.46 | 99 | -0.77 | 47.11 | 149 | -0.83 | 48.95 |
| 50 | -9.44 | 92.74 | 100 | -1.80 | 96.28 | 150 | -1.10 | 68.00 |

**Supplementary Table 2 | Budget sets' parameters, neuroimaging study**

| # | Slope ($p_y/p_x$) | Y intersection | # | Slope ($p_y/p_x$) | Y intersection | # | Slope ($p_y/p_x$) | Y intersection |
|---|---|---|---|---|---|---|---|---|
| 1 | -0.74 | 55.01 | 26 | -1.15 | 70.37 | 51 | -1.00 | 83.80 |
| 2 | -0.23 | 19.60 | 27 | -0.66 | 46.53 | 52 | -0.62 | 44.23 |
| 3 | -0.50 | 42.56 | 28 | -0.11 | 6.07 | 53 | -0.90 | 47.65 |
| 4 | -1.88 | 98.70 | 29 | -0.42 | 21.91 | 54 | -0.89 | 51.79 |
| 5 | -2.54 | 53.16 | 30 | -0.25 | 19.98 | 55 | -7.42 | 87.86 |
| 6 | -1.16 | 97.26 | 31 | -0.46 | 32.99 | 56 | -3.59 | 89.01 |
| 7 | -0.56 | 44.56 | 32 | -1.18 | 93.16 | 57 | -2.35 | 51.47 |
| 8 | -7.96 | 65.39 | 33 | -1.56 | 81.18 | 58 | -0.15 | 10.80 |
| 9 | -1.67 | 95.10 | 34 | -2.31 | 73.22 | 59 | -0.21 | 11.18 |
| 10 | -0.33 | 27.67 | 35 | -0.73 | 56.40 | 60 | -2.84 | 63.98 |
| 11 | -1.38 | 50.11 | 36 | -1.78 | 92.64 | 61 | -0.22 | 18.83 |
| 12 | -1.51 | 56.35 | 37 | -1.54 | 85.23 | 62 | -0.58 | 30.41 |
| 13 | -0.19 | 18.36 | 38 | -2.91 | 82.66 | 63 | -1.11 | 75.59 |
| 14 | -0.17 | 11.35 | 39 | -2.18 | 63.18 | 64 | -1.61 | 55.80 |
| 15 | -1.16 | 99.63 | 40 | -3.65 | 70.37 | 65 | -0.41 | 24.15 |
| 16 | -0.32 | 24.41 | 41 | -0.70 | 46.01 | 66 | -0.10 | 9.13 |
| 17 | -1.24 | 54.26 | 42 | -1.37 | 69.72 | 67 | -0.14 | 8.67 |
| 18 | -0.61 | 44.78 | 43 | -0.35 | 31.17 | 68 | -0.25 | 19.68 |
| 19 | -1.15 | 86.24 | 44 | -0.75 | 60.24 | 69 | -1.60 | 83.55 |
| 20 | -0.21 | 20.88 | 45 | -0.47 | 25.88 | 70 | -1.82 | 96.68 |
| 21 | -0.19 | 15.86 | 46 | -2.59 | 64.18 | 71 | -0.81 | 48.28 |
| 22 | -2.77 | 54.84 | 47 | -1.32 | 58.36 | 72 | -7.75 | 57.36 |
| 23 | -0.59 | 49.72 | 48 | -0.54 | 45.83 | 73 | -3.70 | 82.84 |
| 24 | -1.10 | 89.02 | 49 | -1.64 | 80.46 | 74 | -0.97 | 63.01 |
| 25 | -3.56 | 85.72 | 50 | -9.44 | 92.74 | 75 | -0.63 | 37.93 |

#### Supplementary Table 3 | Mouse features

| # | Name | Description |
|---|---|---|
| **Sample-level features used for pre-processing of the mouse tracking data** | | |
| 1 | OrgTraj | Original mouse trajectory |
| 2 | InterpOrg | Original trajectory interpolated to equal time intervals for 100 samples. |
| 3 | interpTime | Time interpolated to equal intervals for 100 samples. |
| 4 | SR | The interpolated Sampling Rate for 100 samples. |
| 5 | InterpOrgHighRes | Interpolate the original trajectory into an equal time intervals sampling with 1,000 samples per trial (X & Y coordinates) |
| 6 | interpTimeHighRes | The interpolated 1,000 time samples from each trial (in sec) |
| 7 | SRHighRes | The sampling rate of the high-resolution time samples |
| 8 | Angles | The angle of each sample from a straight line to the choice. |
| 9 | Seg_D_Progress | Progress made between two adjacent samples, measured as the distance between one sample to the previous sample. |
| 10 | Comul_Progress | Cumulative progress for each sample |
| 11 | Velocity | Velocity at each sample, measured as the progress between two adjacent samples divided by the time difference between two adjacent samples. |
| 12 | Acceleration | Acceleration at each sample, measured as the difference in velocity between two adjacent sample divided by the time difference between two adjacent samples |
| 13 | Traj2Line | The interpolated trajectory, up to reaching the budget line for the first time. |
| 14 | EqualTraj | The interpolated trajectory up to reaching the budget line, resampled in evenly spaced location, so as to be treated as a curve. |
| 15 | ArcLenCum | The arc length of the curve at each sample |
| 16 | RadiusofCurv | The radius of the curve at each sample |
| 17 | Curvatures | The curvature vector at each sample |
| 18 | D_Predicted | Distance from each sample to the predicted bundle. Predicted bundles are the bundles supposed to be chosen, based on the subject-specific elicited utility parameters |
| **Trial-level features, included in the main analysis** | | |
| 1 | Layover | Time fixated around [0,0], in seconds (latency in movement). |
| 2 | AnglesSTD | Standard deviation of Angles. |
| 3 | MaxProgress | Maximal progress made in a single sample of a trial |
| 4 | Time2MaxProgress | Time passed from beginning of trial to the sample where maximal progress was made. |
| 5 | MedianProgress | Median progress over all samples |
| 6 | VelOver10 | Total time spent in velocities over 10 velocity units (at high speeds) |
| 7 | meanVel | Mean velocity over all samples |
| 8 | MedianVel | Median velocity over all samples. |
| 9 | MaxVel | Maximal velocity over all samples. |
| 10 | MaxVelTime | Time from beginning of trial to the sample of maximal velocity. |
| 11 | meanAcc | Mean acceleration over all samples |
| 12 | MaxAcc | Maximal acceleration over all samples. |
| 13 | EndofTrialVel | Mean velocity at the last 10 samples of a trial. |
| 14 | N_Accs | Count of peaks in acceleration throughout the trial. |
| 15 | ArcLenMax | The maximal Arc length. |
| 16 | Curveness | The sum of the norms of all curvature vectors in the trial |
| 17 | MD | Maximal distance from Choice line |
| 18 | AUC | Area between trajectory and Choice line, only within the budget set |
| 19 | numFixations | Number of individual mouse fixations (mouse stays above **0.2** seconds in the same bin, when bins are **20X20** grids of the trial mouse area (different for each trial). |
| 20 | AveFixTime | Average over all durations of fixations. |
| 21 | MaxFixTime | Maximal duration of fixations. |

| | | |
|---|---|---|
| 22 | MaxFixChoiceDist | Distance from choice coordinates to the coordinates of the longest fixation bin. |
| 23 | numBudgetFixations | Number of fixations near budget line (distance shorter than **2** units) |
| 24 | AboveLine | Time spent above the budget line |
| 25 | XTimeOutofBounds | Time the mouse was out of bounds of the x axis (above 100, below 0) |
| 26 | YTimeOutofBounds | Time the mouse was out of bounds of the y axis (above 100, below 0) |
| 27 | TimeNearPredicted | Time spent in distance under 10 units from the predicted bundle. |
| 28 | TimeBudgetLine | Time spent around the budget line, 2 units above/below it. |
| 29 | XSampEn4 | The sample entropy of the x coordinates, calculated with a window of 4. |
| 30 | YSampEn4 | The sample entropy of the y coordinates, calculated with a window of 4. |
| 31 | Xflips | Numbers of flips on x axis |
| 32 | MaxXflip | Size of the largest flip in X |
| 33 | Yflips | Number of flips on y axis |
| 34 | MaxYflip | Size of the largest flip in Y |

**Supplementary Table 4 | Surviving features from the elastic net analysis, behavioral and neuroimaging studies**

| Behavioral study | | | Neuroimaging study | |
|---|---|---|---|---|
| **Main task** | **Motor task** | **Numerical task** | **Main task** | **Motor task** |
| Xflips*** | Xflips** | Xflips | Xflips | Xflips*** |
| Yflips*** | MaxYflip | Yflips** | Yflips** | MaxYflip |
| MaxXflip*** | MaxProgress | MaxXflip | MaxXflip | MaxProgress |
| MaxYflip*** | meanVel*** | MaxProgress | MaxYflip | medianProgress |
| MaxProgress | MaxVel | medianProgress | MaxProgress | meanVel*** |
| medianProgress | medianVel | Time2MaxProgress | medianProgress | medianVel |
| Time2MaxProgress | VelOver10*** | meanVel | meanVel*** | VelOver10 |
| meanVel*** | EndofTrialVel*** | MaxVel | MaxVel | EndOfTrialVel |
| medianVel | MaxVelTime | medianVel | medianVel | meanAcc |
| VelOver10*** | meanAcc | VelOver10 | VelOver10 | MaxAcc |
| EndofTrialVel | MaxAcc | EndofTrialVel | EndOfTrialVel** | N_Accs |
| MaxVelTime | MaxArcLen | MaxVelTime | MaxVelTime** | MaxArcLen |
| meanAcc | Curvenes | meanAcc | meanAcc | Angle_STD*** |
| MaxAcc | MD*** | MaxAcc | MaxAcc** | MD |
| N_Accs*** | AUC | N_Acc | N_Accs** | AUC** |
| maxArcLen | Layover | MaxArcLen | MaxArcLen | Layover |
| Curveness*** | numFixations | Curvenes** | Curveness | numFixations*** |
| Angle_STD** | MaxFixTime | Angle_STD | Angle_STD | MaxFixChoiceDist |
| MD*** | MaxFixChoiceDist | MD*** | MD*** | AveFixTime*** |
| AUC*** | numBudgetFixations | AUC*** | AUC | AboveLine |
| Layover** | AboveLine | Layover | Layover | XTimeOutofBounds |
| MaxFixTime | XTimeOutofBounds*** | numFixations | numFixations*** | YTimeOutofBounds |
| numBudgetFixations** | TimeBudgetLine | MaxFixTime | MaxFixTime | TimeNearPredicted |
| AveFixTime | XSampEn4*** | MaxFixChoiceDist | MaxFixChoiceDist | TimeBudgetLine |
| AboveLine*** | YSampEn4 | numBudgetFixations | AboveLine | YSampEn4 |
| YTimeOutofBounds | | AveFixTime | XTimeOutofBounds*** | |
| TimeNearPredicted*** | | AboveLine | YTimeOutofBounds** | |
| TimeBudgetLine*** | | XTimeOutofBounds*** | TimeNearPredicted** | |
| XSampEn4*** | | YTimeOutofBounds | TimeBudgetLine | |
| YSampEn4*** | | TimeNearPredicted*** | XSampEn4 | |
| | | TimeBudgetLine** | | |
| | | XSampEn4*** | | |
| | | YSampEn4 | | |

(**) $p<0.05$, (***) $p<0.01$, significantly correlated with MMI in a multiple linear model with subjects FEs (random intercept).

**Supplementary Table 5 | Surviving features from the elastic net analysis, replication study**

| Replication study | | |
|---|---|---|
| **Main task** | **Motor task** | **Numerical task** |
| Xflips | Xflips | Xflips*** |
| Yflips*** | Yflips | Yflips |
| MaxXflip*** | MaxXflip | MaxXflip |
| MaxYflip*** | MaxProgress | meanVel |
| MaxProgress | meanVel | MaxVel*** |
| meanVel*** | MaxVel | VelOver10 |
| MaxVel** | medianVel** | Curvenes |
| medianVel*** | VelOver10*** | Angle_STD*** |
| VelOver10** | EndofTrialVel | MD*** |
| EndofTrialVel | MaxVelTime | AUC*** |
| meanAcc | N_Acc | Layover*** |
| MaxAcc | MaxArcLen*** | AveFixTime*** |
| N_Accs** | Curvenes** | AboveLine |
| maxArcLen | Angle_STD*** | YTimeOutofBounds |
| Curveness** | MD*** | XSampEn4 |
| Angle_STD*** | AUC | YSampEn |
| MD*** | numFixations | |
| AUC*** | MaxFixTime** | |
| numFixations*** | MaxFixChoiceDist | |
| MaxFixTime*** | numBudgetFixations | |
| MaxFixChoiceDist*** | AboveLine | |
| numBudgetFixations | XTimeOutofBounds** | |
| AveFixTime | YTimeOutofBounds** | |
| AboveLine** | TimeNearPredicted | |
| XTimeOutofBounds*** | TimeBudgetLine | |
| TimeNearPredicted*** | XSampEn4 | |
| TimeBudgetLine*** | | |
| XSampEn4*** | | |
| YSampEn4** | | |

(**) $p<0.05$, (***) $p<0.01$, significantly correlated with MMI in a multiple linear model with subjects FEs (random intercept).

#### Supplementary Table 6 | Psychophysiological Interaction analysis results

| | **Seed** | | | |
|---|---|---|---|---|
| ***M1*** | **vmPFC** | **vStr** | **dACC** | **PCC** |
| Intercept | -0.002<br>(0.004) | -0.004<br>(0.004) | -0.007<br>(0.004) | -0.003<br>(0.004) |
| RT | -0.399***<br>(0.009) | -0.124***<br>(0.009) | -0.258<br>(0.009)*** | -0.332***<br>(0.009) |
| Slope | 0.008<br>(0.004) | 0.012**<br>(0.005) | -0.003<br>(0.005) | 0.005<br>(0.005) |
| Endowment | -0.045***<br>(0.007) | -0.017*<br>(0.007) | -0.013<br>(0.007) | -0.009<br>(0.008) |
| MMI | 0.009<br>(0.005) | -0.007<br>(0.005) | -0.002<br>(0.005) | -0.002<br>(0.005) |
| M1 | 0.364***<br>(0.005) | 0.525***<br>(0.005) | 0.536***<br>(0.005) | 0.495***<br>(0.005) |
| MMI×M1 | 0.013*<br>(0.005) | **0.012***<br>(0.005)** | **0.036***<br>(0.005)** | **0.019***<br>(0.005)** |
| ***SMA*** | **vmPFC** | **vStr** | **dACC** | **PCC** |
| Intercept | -0.001<br>(0.004) | 0.001<br>(0.004) | -0.002<br>(0.003) | -0.001<br>(0.004) |
| RT | -0.364***<br>(0.008) | -0.055***<br>(0.008) | -0.251***<br>(0.007) | -0.244***<br>(0.008) |
| Slope | 0.008<br>(0.005) | 0.012**<br>(0.004) | -0.003<br>(0.004) | 0.005<br>(0.004) |
| Endowment | -0.062***<br>(0.007) | -0.039***<br>(0.007) | -0.038***<br>(0.006) | -0.029***<br>(0.007) |
| MMI | 0.010*<br>(0.005) | 0.000<br>(0.004) | 0.006<br>(0.004) | 0.000<br>(0.005) |
| M1 | 0.453***<br>(0.004) | 0.608***<br>(0.004) | 0.765***<br>(0.003) | 0.523***<br>(0.004) |
| MMI×SMA | 0.004<br>(0.004) | -0.004<br>(0.004) | **0.012***<br>(0.003)** | 0.007<br>(0.004) |

(*) p<0.05, (**) p<0.01, (***) p<0.001. Standard errors are in parentheses. PPI analysis results for all value regions seeds with M1 and SMA as regressors. Significant PPI terms are marked in bold. Note that the vmPFC-M1 PPI was not significant after using Bonferroni correction for multiple comparisons.

**Supplementary Table 7 | Adj $R^2$ from a regression that only includes the "winning features" (MD and meanVel)**

| | Main task | Motor task | Numerical task |
|---|---|---|---|
| **Behavioral study** | 1.9% | 0.5% | 1.7% |
| **Neuroimaging study** | 2% | 0.8% | |
| **Replication study** | 0.6% | 2.7% | 3.0% |
